# Tuning viral virulence and cytokine release enforces oncotropism and eliminates systemic toxicity: A framework for next-generation oncolytics

**DOI:** 10.64898/2026.11.20.768167

**Authors:** Reza Rezaei, Gabriel St-Laurent, Nika Koshiki Zamani, Anya Alsamaraie, Zumama Khalid, Zacharie Drouin, Zahra Ameli, Phuong Anh Nguyen, Mahsa Ahmadi, Maria Fernanda Solorza Reyes, Pedram Abdollahpour, Georgia L. Cheung, Arezoo Sadeghipour, Julia Petryk, Fazel Mohammadi, Miles Da Silva, Harkirat Dhillon, Kian Jafari, Samuel Workenhe, Raymund Wellinger, Damian Carragher, Marilyne Labrie, Shervin Taslimi, John C. Bell, Taha Azad

## Abstract

Oncolytic virotherapy has long balanced safety and potency through permanent attenuation and, increasingly, constitutive inflammatory payloads. Yet attenuation can restrict replication in heterogeneous tumors, whereas unregulated cytokine expression can cause systemic toxicity and promote premature viral clearance. Here, we establish an alternative vaccinia framework that replaces fixed attenuation with reversible chemogenetic control of the essential replication gene E1L and coordinates replication with graded interleukin-12 (IL-12) expression through a hierarchical, drug-gated circuit. E1L control enabled inducible replication with no detectable consensus-level escape under tested selection conditions. Following intravenous administration, the fully induced virus maintained high intratumoral titers while reducing infectious-virus recovery across peripheral organs by orders of magnitude relative to thymidine-kinase-deleted vaccinia and the induced parent vector, without marked weight loss. Across syngeneic tumor models, dual induction prolonged survival and was associated with cytotoxic CD8+ T-cell and inflammatory macrophage states. This architecture establishes post-entry functional oncotropism through control of productive viral persistence.

## INTRODUCTION

Oncolytic virotherapy emerged from clinical observations of tumor regressions following natural viral infections, establishing a therapeutic strategy that couples direct oncolysis with antitumor immune activation^1,2^. The field’s central design challenge lies in maximizing viral replication and spread within tumors while strictly restraining virulence in healthy tissue^3^. To establish a safety margin, conventional engineering frameworks have relied on permanent gene deletions targeting virulence-associated genes, such as vaccinia virus thymidine kinase^4^, herpes simplex virus 1 (HSV-1) ICP34.5^5^, E1B-55K in adenovirus^6^, or attenuation of the matrix protein in vesicular stomatitis virus (VSV)^7^. However, this strategy poses a functional risk: permanently removing viral functions that facilitate host tissue spread can compromise the replicative fitness required to overcome solid tumor barriers^5–8^. Preclinical HSV-1 studies highlight this fitness cost: restoring tumor-restricted ICP34.5 expression substantially enhances viral spread and therapeutic efficacy in glioma^5^, while preserving even a single copy of ICP34.5 enables NV1020 to outperform the more heavily deleted G207 vector in advanced tumor models^8^. Clinical experience with the thymidine kinase-positive vaccinia strain ACAM2000 provides a complementary perspective: in a Phase I study using autologous stromal vascular fraction cells for delivery, some patients had viral DNA detectable in blood for weeks without serious treatment-related toxicity^9^. These findings motivate an alternative design principle: rather than permanently deleting viral machinery, placing essential replication genes under reversible post-administration control provides dynamic tuning of virulence while preserving the capacity for robust oncolytic propagation^10^.

However, regulating viral replication kinetics alone does not inherently enforce spatial oncotropism; systemically administered conditionally replicating vectors can still seed permissive healthy tissues^3^. While structural envelope engineering has been explored to retarget viral platforms such as HSV-1^11,12^, vaccinia presents a distinct engineering challenge because multiple viral proteins contribute to attachment and entry, uptake pathways vary by strain and cell type, and no single canonical entry receptor has been established^13,14^. Rather than structural envelope modification, we hypothesized that equipping oncolytic vectors with a potent pro-inflammatory cytokine could establish an immune-mediated mechanism of oncotropism, alerting host defenses to accelerate viral clearance in normal organs while sustaining replication within the immunosuppressive tumor microenvironment^15^. Interleukin-12 (IL-12) represents a well-suited candidate, as it coordinates both innate and adaptive immunity by stimulating natural killer and cytotoxic T cell activity, repolarizing tumor-associated macrophages toward an inflammatory phenotype, and driving robust interferon-gamma production^16^. Because the clinical development of recombinant IL-12 has been severely compromised by systemic toxicity, coupling its expression directly to a switch-regulated oncolytic platform offers a dual therapeutic rationale: promoting tumor-biased viral persistence through localized immune alerting, while limiting systemic cytokine exposure to reduce the risk of dose-limiting pathology^10,17^.

Here, we engineered a conditionally replicating vaccinia virus platform (CRVV) by placing the essential viral gene E1L under doxycycline-inducible control and coupling it to a coordinated, small-molecule-regulated circuit driving interleukin-12 expression (CRVV12). Following systemic intravenous administration in immunocompromised mice, this platform demonstrated marked tumor-biased persistence and superior tolerability relative to a benchmark thymidine kinase-deleted (ΔTK) vector, maintaining high infectious titers within the tumor bed while substantially reducing viral burden in healthy organs without marked weight loss during the study. As a monotherapy, dual-switch induction suppressed tumor growth and extended survival in several syngeneic tumor models. Immune profiling showed enrichment of cytotoxic CD8+ T cell and inflammatory macrophage states in treated tumors, and checkpoint blockade further improved antitumor activity in selected models. Together, these findings demonstrate that coordinated chemogenetic control provides a viable framework for tuning viral virulence and tumor-biased activity while preserving robust intratumoral replication.

## RESULTS

### Inducible control of E1L gates vaccinia replication without detectable mutational escape after inducer-free growth

Chemogenetic control of essential viral genes has previously been used to impose complete replication arrest for single-cycle vaccine applications^10,18,19^. Oncolytic development presents a distinct engineering requirement: the regulated locus must preserve sufficient replication for therapeutic propagation while restricting genetic escape under selective conditions that favor the outgrowth of replication-competent variants. To identify a suitable regulatory node, we targeted three essential vaccinia genes operating at distinct stages of the viral lifecycle: D13L, which mediates virion membrane morphogenesis^18^; F17R, which contributes to viral core assembly^20^; and E1L, which encodes the catalytic subunit of the viral poly(A) polymerase required for mRNA processing^21^. Each locus was engineered under a synthetic TetO-containing promoter repressed by constitutively expressed TetR (Fig. 1A). We then assessed plaque formation in U2OS cells in the presence or absence of doxycycline (Fig. 1B, C). Regulation of F17R yielded no measurable doxycycline-dependent change in plaque expansion, whereas regulation of D13L imposed near-complete repression without inducer and supported robust plaque expansion upon doxycycline addition. In contrast, regulation of E1L produced an intermediate phenotype, retaining limited plaque formation in the uninduced state while driving substantially greater viral spread upon induction (Fig. 1B, C). Multistep growth kinetics in HeLa cells confirmed that uninduced E1L-regulated virus, designated CRVV, remained replication-competent but yielded viral titers approximately one to two orders of magnitude below wild-type vaccinia virus and vaccinia virus with thymidine kinase deletion (VV-ΔTK) at late time points. Doxycycline induction restored CRVV yields to the range of replication-competent controls by 48 hours (Fig. S1A). This inducible phenotype was preserved across HEK293, HeLa, and Vero lines, where doxycycline enhanced viral spread and cytolysis relative to uninduced conditions (Fig. S1B, C).

**Fig. 1.**
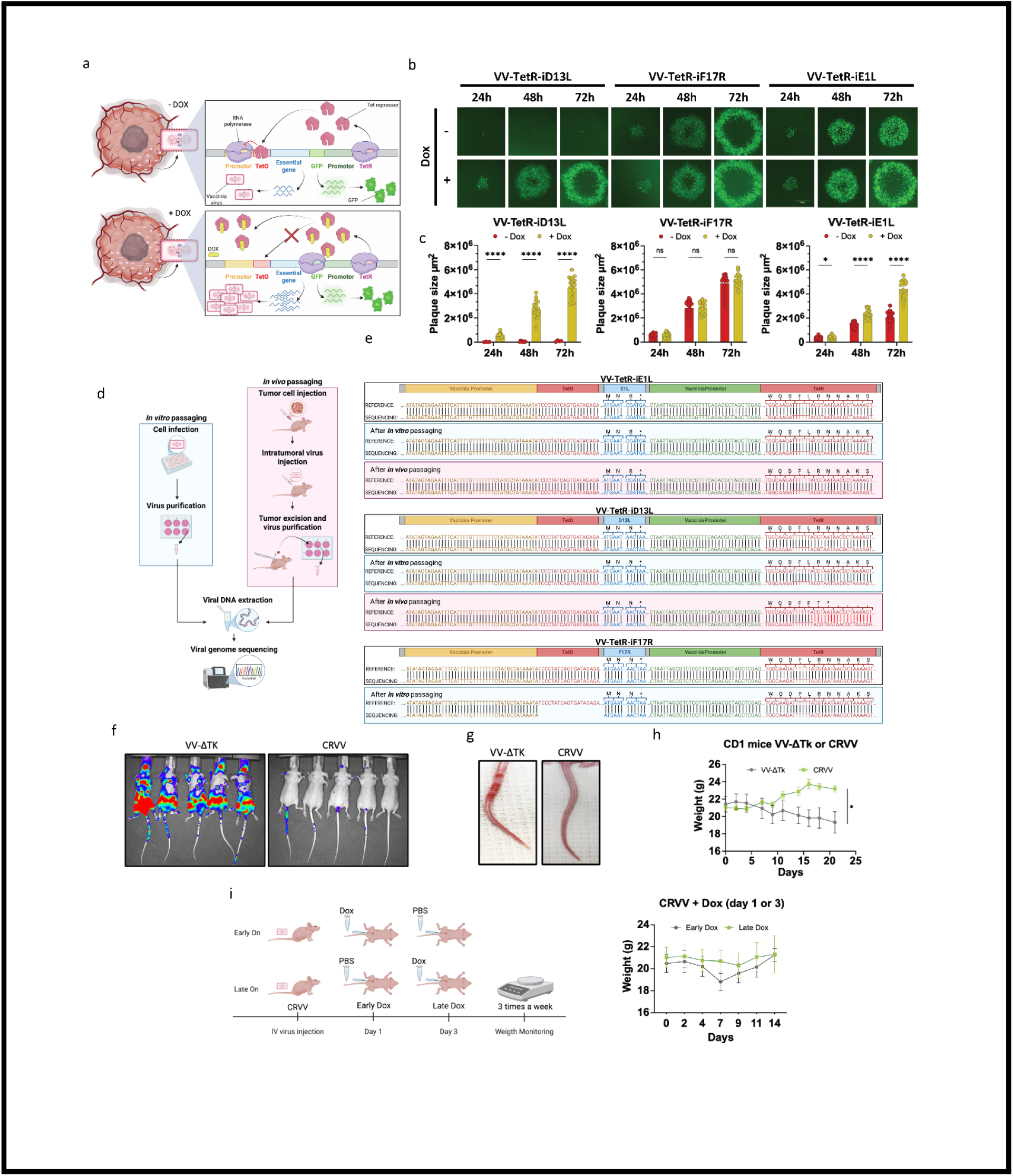
Locus-dependent selective pressure drives mutational escape under complete shutoff, establishing E1L as a stable oncolytic replication switch. (A) Schematic representation of the chemogenetic replication switch, showing an essential viral gene regulated by the TetO/TetR system in the absence (-Dox) or presence (+Dox) of doxycycline. (B) Representative fluorescence microscopy images of viral plaques in U2OS cells infected with VV-TetR-iD13L, VV-TetR-iF17R, or VV-TetR-iE1L (MOI 0.00001) with or without doxycycline (100 ng/mL) in the media at 24, 48, and 72 h post-infection. (C) Quantification of viral plaque surface area (µm²) in U2OS cells infected with VV-TetR-iD13L, VV-TetR-iF17R, or VV-TetR-iE1L under -Dox and +Dox conditions at 24, 48, and 72 h post-infection. (D) Schematic of the inducer-free escape assays. For the in vitro assay, U2OS cells were infected once without doxycycline, followed by isolation and deep sequencing of individual large plaques. For the in vivo assay, viruses were administered intratumorally to CD-1 nude mice bearing subcutaneous HT-29 xenografts without doxycycline. Virus recovered from disrupted tumors was amplified in cell culture and used for the subsequent intratumoral inoculation for a total of three sequential rounds, followed by plaque isolation and deep sequencing. (E) Alignment of viral genome sequencing reads covering the promoter-TetO regulatory regions of VV-TetR-iE1L, VV-TetR-iD13L, and VV-TetR-iF17R following in vitro and in vivo passaging relative to reference sequences. (F) In vivo bioluminescence imaging (IVIS) of CD1 nude mice 48 h post-intravenous injection with 1 × 10^7 PFU of VV-ΔTK or CRVV. (G) Representative tail photographs of CD1 nude mice taken at day 21 post-intravenous injection with VV-ΔTK or CRVV. (H) Longitudinal body weight tracking of CD1 nude mice following intravenous injection of VV-ΔTK or CRVV (1 × 10^7 PFU). (I) Dosing timeline and longitudinal body weight monitoring of CD1 nude mice intravenously injected with 1 × 10^7 PFU of CRVV followed by doxycycline administration (625 mg/kg diet) starting on day 1 (Early Dox) or day 3 (Late Dox) post-infection. In vitro data in (C) represent mean +/- SEM from n = 3 replicates. In vivo data in (H) and (I) represent mean +/- SEM from n = 5 mice per group. P values in (C) were determined by two-way ANOVA with Tukey’s post-hoc test. P < 0.01, ****P < 0.0001, ns, not significant..

To assess genetic stability under selective pressure, the regulated vectors underwent an in vitro plaque-outgrowth screen and three rounds of serial intratumoral passage in HT-29 xenografts, followed by plaque isolation and deep sequencing (Fig. 1D). Consensus analysis of the resulting sequencing reads demonstrated locus-dependent genetic escape. The F17R-regulated vector lost TetO sequences during in vitro propagation, whereas the D13L-regulated vector recovered after in vivo selection acquired a multi-adenine insertion within TetR, generating a frameshift predicted to truncate the repressor. In contrast, no consensus-level sequence alterations were detected within the interrogated E1L-regulated cassette (CRVV) following either the single-round in vitro screen or the three-round serial in vivo selection (Fig. 1E).

We next evaluated whether the attenuated basal replication of uninduced CRVV was sufficient to limit systemic viral activity and associated morbidity in an immunocompromised host. CD-1 nude mice received intravenous injections of either uninduced CRVV or the benchmark VV-ΔTK vector. At 48 hours post-injection, VV-ΔTK produced widespread bioluminescent reporter signal, whereas CRVV generated markedly reduced signal throughout the animal (Fig. 1F). Over a 21-day observation period, VV-ΔTK-treated mice developed visible tail pox lesions (Fig. 1G) accompanied by progressive body weight loss (Fig. 1H). Conversely, mice receiving uninduced CRVV maintained stable body weight and remained free of visible tail lesions throughout the study (Fig. 1G, H). Finally, to determine whether the timing of replication induction influenced host tolerability, mice administered with intravenous CRVV initiated doxycycline treatment on either day 1 or day 3 post-infection (Fig. 1I). Induction beginning on day 1 caused a transient reduction in body weight followed by steady recovery, whereas delaying induction to day 3 resulted in minimal weight deviation over the observation window (Fig. 1I). Together, these findings identify E1L as an optimal regulatory switch that couples meaningful basal restriction with inducible restoration of replication, exhibiting no detectable consensus-level escape under in vitro and serial in vivo selection while substantially mitigating systemic viral activity and morbidity.

### Hierarchical dual-input gating coordinates vaccinia replication with graded interleukin-12 output

We first evaluated the oncolytic efficacy and immune remodeling induced by uncoupled CRVV monotherapy across syngeneic tumor models. In subcutaneous B16-F10 melanoma, intratumoral administration of CRVV under uninduced (-Dox) or induced (+Dox) conditions showed tumor-associated bioluminescent reporter activity under both states at day 5 post-injection (Fig. S2A), with both uninduced and induced vectors slowing tumor growth kinetics without a significant separation between them (Fig. S2B). Flow cytometric profiling of the tumor microenvironment showed a higher proportion of CD3+ T cells and a lower proportion of CD11b+ myeloid cells within the CD45+ compartment in both CRVV treatment groups relative to PBS controls, without significant compositional differences between the uninduced and induced states (Fig. S2C-E). In subcutaneous CT26-LacZ colorectal tumors, both uninduced and induced CRVV suppressed tumor growth and prolonged survival, again without a clear separation between the two states (Fig. S2F, G). In contrast, in the more aggressive intraperitoneal MC38 tumor challenge, induced CRVV (+Dox) significantly extended animal survival, whereas animals treated with PBS, VV-ΔTK with or without doxycycline, or uninduced CRVV reached survival endpoints within 25 days post-inoculation (Fig. 2A). In a separate experiment, profiling of the immune compartment in subcutaneous MC38 tumors revealed a distinct phenotypic shift among CD4+ T cells, where representative plots showed a larger CD62L+ fraction after CRVV treatment relative to VV-ΔTK, while CD8+ T cells maintained a predominantly CD62L-CD44+ phenotype across groups (Fig. 2B). Systemically, total splenocyte counts trended higher and PMN-MDSC frequencies in the spleen and tumor trended lower than those observed with VV-ΔTK (Fig. S2H, I), while CRVV significantly increased splenic T-cell frequency (Fig. S2J). Frequencies of PD-1+, TIGIT+, LAG-3+, and TIM-3+ CD4+ T cells trended lower in CRVV-treated mice, whereas frequencies of checkpoint-positive CD8+ T cells showed no consistent treatment-associated pattern (Fig. S2K, L). Together, these studies showed that E1L induction enhanced therapeutic activity in a model-dependent manner but did not consistently produce durable tumor control, motivating incorporation of a regulated immunostimulatory payload.

**Fig. 2.**
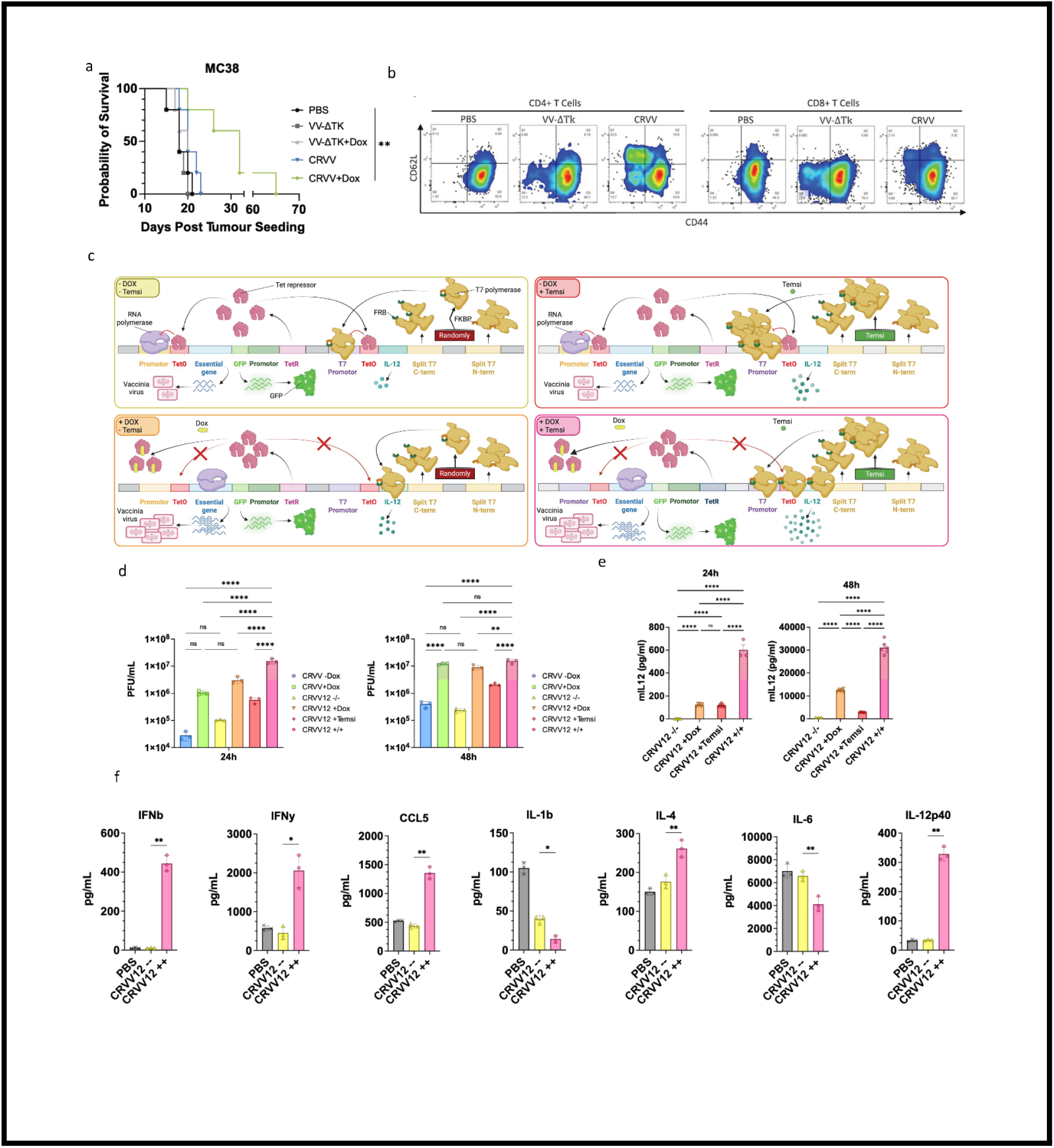
Dual-switch chemogenetic gating couples essential gene expression to split-T7 activation, limiting basal output and enabling graded interleukin-12 induction. (A) Kaplan-Meier survival curves of C57BL/6 mice bearing intraperitoneal MC38 tumors treated with intraperitoneal injections of PBS, VV-ΔTK, VV-ΔTK + Dox, CRVV, or CRVV + Dox (1 × 10^7 PFU per dose) on days 7, 9, and 11 post-tumor inoculation. (B) Representative flow cytometry plots of CD62L and CD44 surface expression on CD4+ (left) and CD8+ (right) T cells isolated from subcutaneous MC38 tumors 6 days post-intratumoral injection of PBS, VV-ΔTK, or CRVV (1 × 10^7 PFU). (C) Architectural schematic of the CRVV12 dual-switch circuit. Viral replication is controlled by TetR-mediated repression of the essential gene E1L. Expression of murine IL-12 is controlled by a T7 promoter followed by a tetracycline operator (TetO), transcribed by a split-T7 RNA polymerase that reconstitutes upon temsirolimus-mediated heterodimerization. (D) Multi-step viral growth kinetics in HeLa cells infected with CRVV or CRVV12 (MOI 0.01) at 24 and 48 h post-infection in the absence or presence of doxycycline (100 ng/mL), temsirolimus (10 nM), or both. Total viral progeny were titered on U2OS cells. (E) ELISA quantification of murine IL-12 (pg/mL) in supernatants of U2OS cells infected with CRVV12 (MOI 0.01) at 24 and 48 h post-infection across the indicated drug combinations. (F) Absolute concentrations (pg/mL) of IFN-β, IFN-γ, CCL5, IL-1β, IL-4, IL-6, and IL-12p40 released from naive Balb/c splenocytes co-cultured for 24 h with filtered conditioned media from MC38 cells infected with CRVV12 (MOI 0.001) for 24 h under uninduced (CRVV12 -/-) or fully induced (CRVV12 +/+) conditions relative to PBS controls. In vitro data in (D) to (F) represent mean +/- SEM from n = 3 replicates. In vivo survival data in (A) represent n = 5 mice per group. P values in (A) were determined by log-rank (Mantel-Cox) test; P values in (D) to (F) were determined by one-way or two-way ANOVA with Tukey’s post-hoc test. *P < 0.05, P < 0.01, ***P < 0.001, ****P < 0.0001, ns, not significant.

To achieve coordinated control over potent therapeutic payloads, we integrated a second chemogenetic layer into the CRVV backbone to regulate interleukin-12 expression (CRVV12). In this design, the shared TetR node simultaneously represses both the essential viral gene E1L and a TetO-containing T7 promoter driving murine IL-12, while payload transcription additionally requires reconstitution of split-T7 RNA polymerase through temsirolimus-induced FKBP-FRB heterodimerization^10^ (Fig. 2C). The circuit was designed to limit basal payload expression through TetR-mediated transcriptional repression and the reduced viral template abundance associated with the uninduced replication state, establishing a stepwise gradient of IL-12 induction upon switch activation (Fig. 2C). Multistep viral growth kinetics in HeLa cells confirmed that CRVV12 replication remained doxycycline responsive: doxycycline-containing conditions increased viral yield relative to the corresponding uninduced states at both 24 and 48 hours, whereas temsirolimus alone did not restore replication to the levels reached under doxycycline-containing conditions (Fig. 2D). Western blotting confirmed minimal IL-12 expression in the uninduced state and drug-responsive expression following single- or dual-input activation (Fig. S3A, B). In culture supernatants, ELISA quantification demonstrated negligible IL-12 secretion under uninduced conditions (-/-), intermediate release following single-agent activation (+Dox or +Temsi), and a pronounced increase under dual-switch induction (+/+), reaching approximately 600 pg/mL at 24 hours and exceeding 30,000 pg/mL by 48 hours post-infection (Fig. 2E). Exposing naive splenocytes to conditioned medium from infected MC38 cells produced a distinct cytokine profile. Relative to the PBS and uninduced CRVV12 conditions, conditioned medium from fully induced CRVV12 (+/+) was associated with higher concentrations of IFN-γ, IFN-β, CCL5, IL-4, and IL-12p40 and lower concentrations of IL-1β and IL-6 (Fig. 2F), together with broader modulation of additional cytokines and chemokines (Fig. S3C). Together, these results establish CRVV12 as a hierarchical regulatory architecture that maintains doxycycline-responsive viral replication while producing low basal, intermediate single-input, and high dual-input IL-12 expression.

### Coordinated viral replication and interleukin-12 expression establish tumor-biased persistence and mitigate systemic morbidity

To evaluate whether payload coupling altered vector persistence and tolerability, we compared uncoupled CRVV with payload-coupled CRVV12 following intratumoral administration in immunodeficient CD-1 nude mice bearing subcutaneous HT-29 colorectal xenografts (Fig. 3A). Following intratumoral injection, mice receiving uncoupled CRVV under doxycycline-mediated replication induction (CRVV + Dox) exhibited progressive body weight loss exceeding 15% and marked treatment-associated mortality over the observation period (Fig. 3B-D). In contrast, mice receiving CRVV12 under dual induction (CRVV12 +/+) or under uninduced and temsirolimus-only conditions maintained stable body weight with 100% overall survival through day 35, while CRVV12 with doxycycline alone maintained approximately 75% survival (Fig. 3C, D). Longitudinal bioluminescence imaging showed that while uncoupled CRVV produced persistent and increasingly disseminated bioluminescent reporter signal, CRVV12 reporter signal remained predominantly associated with the tumor region (Fig. 3B).

**Fig. 3.**
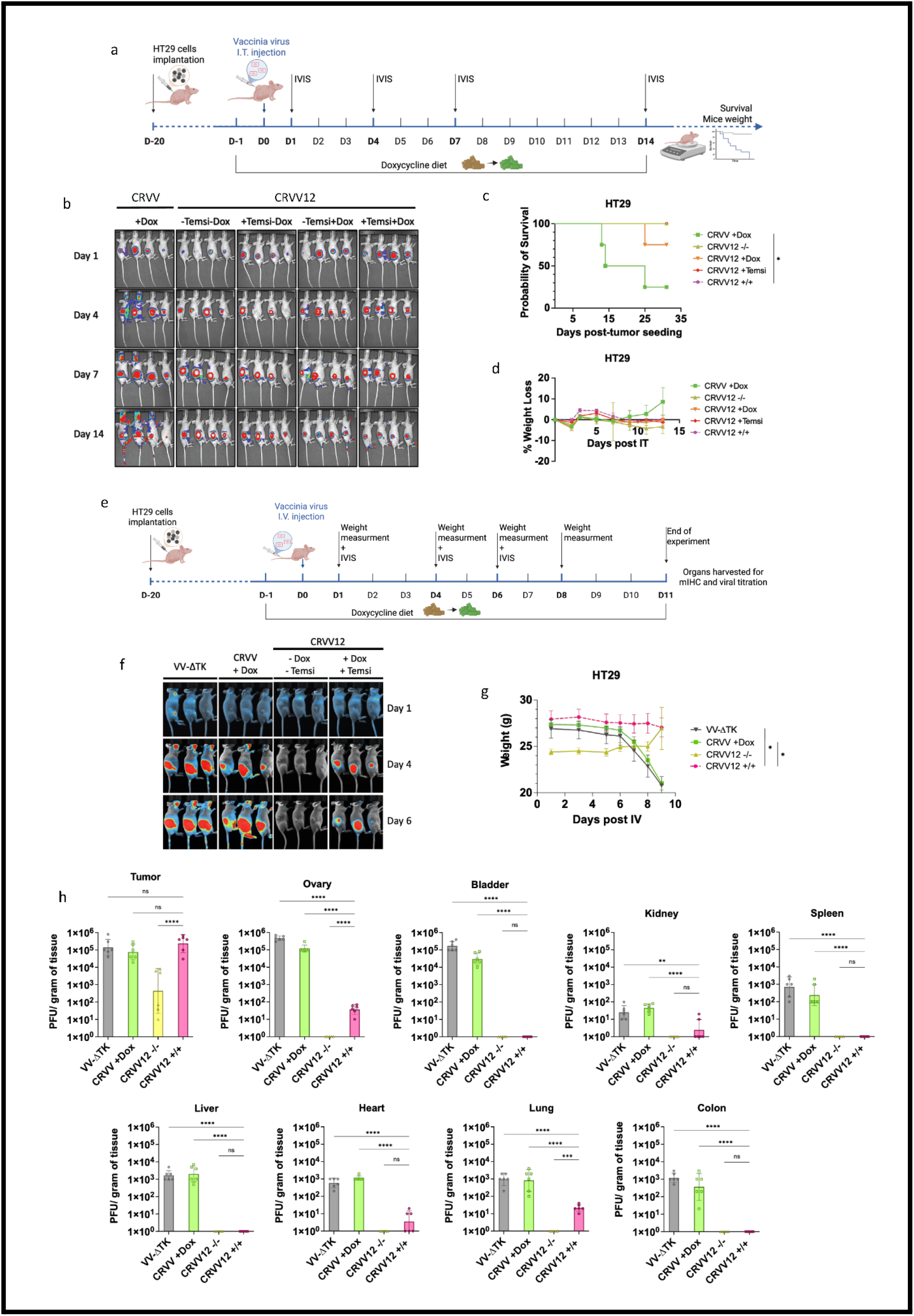
Localized interleukin-12 expression confines viral replication to the tumor bed, preventing systemic dissemination and fatal off-target toxicity. (A) Schematic representation of the experimental timeline for intratumoral (I.T.) viral administration in CD1 nude mice bearing subcutaneous HT29 human colorectal adenocarcinoma xenografts. Mice were maintained on control or doxycycline diet (625 mg/kg) and received intraperitoneal injections of temsirolimus (2.5 mg/kg) as indicated. (B) Representative longitudinal in vivo bioluminescence imaging (IVIS) of CD1 nude mice at days 1, 4, and 6 following I.T. injection of CRVV or CRVV12 (1 × 10^7 PFU). (C) Kaplan-Meier survival curves of CD1 nude mice bearing subcutaneous HT29 tumors following I.T. administration of CRVV or CRVV12 across indicated drug conditions. (D) Percentage body weight change in CD1 nude mice tracked through day 14 post-I.T. viral administration. (E) Schematic overview of the systemic intravenous (I.V.) delivery study design in CD1 nude mice bearing subcutaneous HT29 tumors. Mice received an I.V. injection of 1 × 10^7 PFU of VV-ΔTK, CRVV, or CRVV12 on day 0 with or without doxycycline diet and intraperitoneal temsirolimus administration (days -1, 1, and 4), followed by tissue harvesting on day 11. (F) Representative IVIS bioluminescence images of CD1 nude mice on days 1, 4, and 6 following I.V. injection of VV-ΔTK, CRVV, or CRVV12 under indicated treatment regimens. (G) Longitudinal body weight tracking of CD1 nude mice following I.V. administration of VV-ΔTK, CRVV, or CRVV12. (H) Viral biodistribution across tissues at day 11 post-I.V. injection. Viral load (PFU/gram of tissue) was quantified by plaque assay on U2OS cells from harvested tumor, ovary, bladder, kidney, spleen, liver, heart, lung, and colon. In vivo data in (C), (D), (G), and (H) represent mean +/- SEM from n = 5 mice per group. P values in (C) were determined by log-rank (Mantel-Cox) test; P values in (D), (G), and (H) were determined by one-way or two-way ANOVA with Tukey’s post-hoc test. *P < 0.05, **P < 0.01, ***P < 0.001, ****P < 0.0001, ns, not significant.

We next evaluated whether payload coupling altered viral persistence and tissue distribution following systemic intravenous administration (Fig. 3E). Whole-body bioluminescence imaging of CD-1 nude mice intravenously injected with control VV-ΔTK or induced CRVV (+Dox) revealed diffuse extra-tumoral reporter signal by days 4 and 6 post-infusion, accompanied by marked body weight loss (Fig. 3F, G). In contrast, mice receiving fully induced CRVV12 (+/+) displayed reporter signal predominantly localized to the subcutaneous tumor, maintaining stable body weight throughout the study (Fig. 3F, G). Quantitative viral titration at day 11 confirmed that CRVV12 (+/+) retained high intratumoral viral titers exceeding 10^5 PFU/g of tumor tissue, while sharply reducing infectious virus recovery across normal organs relative to VV-ΔTK and induced CRVV (Fig. 3H). Specifically, while VV-ΔTK and induced CRVV yielded high viral loads across peripheral organs, infectious CRVV12 (+/+) was below the limit of detection in the bladder, spleen, liver, and colon, and was reduced by several orders of magnitude in the ovary, kidney, heart, and lung (Fig. 3H). Together, these findings show that coupling inducible viral replication to regulated IL-12 expression shifts infectious virus persistence toward the tumor bed while mitigating systemic morbidity, supporting coordinated virulence and payload gating as an effective strategy for functional oncotropism.

### Single-cell profiling reveals cytotoxic lymphoid activation and inflammatory myeloid remodeling following dual-switch CRVV12 induction

We evaluated the therapeutic activity of CRVV12 in the subcutaneous B16-F10 melanoma model (Fig. 4A and Fig. S4A). Dual-switch induction (CRVV12 +/+) delayed tumor outgrowth and significantly extended overall survival compared to control cohorts (Fig. 4A and Fig. S4A). Flow-cytometric profiling of tumor-infiltrating leukocytes revealed distinct treatment-associated shifts in the composition of the CD45+ compartment (Fig. 4B). Quantification demonstrated that this antitumor response occurred without an increase in overall lymphocyte representation: fully induced tumors displayed lower relative frequencies of CD8+ T cells, CD4+ T cells, and NK cells alongside a higher relative frequency of macrophages within the CD45+ infiltrate compared to uninduced controls (Fig. S4B). To resolve the cellular states associated with this compositional profile at single-cell resolution, we performed scRNA-seq on CD45+ and CD45-fractions pooled from five tumors per condition and recombined at a 1:1 ratio for sequencing (Fig. 4C and Fig. S5A-D). Consistent with the cytometric findings, global clustering resolved major cell lineages and showed a reduction in relative lymphoid representation together with increased relative representation of the myeloid compartment in the pooled CRVV12 (+/+) condition (Fig. 4C, Fig. S5A-B).

**Fig. 4.**
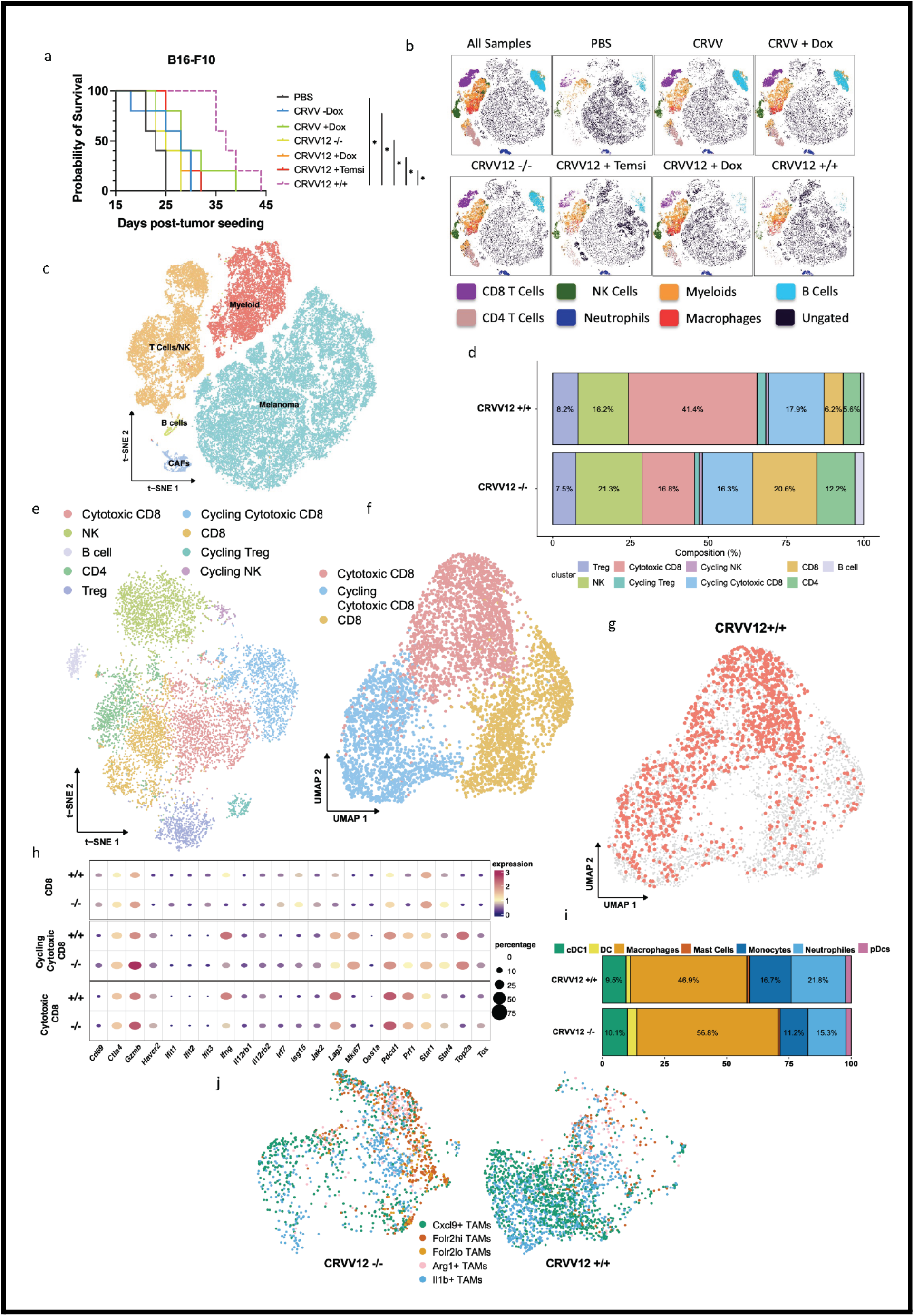
Dual-switch CRVV12 induction remodels cytotoxic T-cell and macrophage states in melanoma. (A) Kaplan-Meier survival curves of C57BL/6 mice bearing subcutaneous B16-F10 melanoma tumors treated with intratumoral injections of PBS, CRVV -Dox, CRVV +Dox, CRVV12 -/-, CRVV12 +Dox, CRVV12 +Temsi, or CRVV12 +/+ (1 × 10^7 PFU per dose) on days 7, 9, and 11 post-tumor inoculation. (B) Representative t-SNE maps from flow cytometry profiling of CD45+ tumor-infiltrating immune cells harvested 4 days post-infection across indicated treatment groups. (C) t-SNE projection of total single-cell transcriptomes from B16-F10 tumors harvested on day 4 post-treatment (n = 5 biological replicates pooled, FACS-sorted for CD45+ and CD45-cells and re-pooled at a 1:1 ratio into two technical sequencing replicates). (D) Stacked bar chart showing the relative composition (%) of tumor-infiltrating lymphoid subclusters in CRVV12 +/+ versus CRVV12 -/- tumors. (E) t-SNE visualization of the tumor-infiltrating lymphoid compartment annotated by functional cell subtype. (F) UMAP projection of the CD8+ T cell lineage. (G) UMAP projection of CD8+ T cells highlighting cells originating from the CRVV12 +/+ condition. (H) Dot plot showing normalized expression and percentage of expressing cells for key effector, activation, cytokine receptor, and exhaustion markers across CD8+ T cell subsets in CRVV12 -/- and CRVV12 +/+ tumors. (I) Stacked bar chart showing the composition (%) of tumor-associated myeloid lineages in CRVV12 +/+ versus CRVV12 -/- tumors. (J) t-SNE projections of tumor-associated macrophages (TAMs) in CRVV12 -/- and CRVV12 +/+ tumors. In vivo survival data in (A) represent n = 5 mice per group; P values were determined by log-rank (Mantel-Cox) test .

We next resolved heterogeneity within tumor-infiltrating lymphocytes by subclustering the lymphoid compartment (Fig. 4D, E). Transcriptional profiling separated CD8+ T cells into three discrete states: a basal CD8 cluster, a cytotoxic CD8 cluster, and a cycling cytotoxic CD8 cluster (Fig. 4D, E). Dimensionality reduction by UMAP revealed a clear difference in the distribution of CD8+ T-cell states between treatment conditions: whereas uninduced CRVV12 (-/-) tumors were enriched for the basal CD8 cluster, cells from fully induced CRVV12 (+/+) tumors concentrated predominantly within the cytotoxic CD8 cluster, while the cycling cytotoxic fraction remained comparable between cohorts (Fig. 4F, G). Within this population, cytotoxic effectors including Gzmb and Prf1 were maintained at broadly comparable levels across conditions, whereas Ifng expression was higher in CRVV12 (+/+) tumors (Fig. 4H). Pseudotime analysis was consistent with this phenotypic skewing, positioning cells from fully induced tumors preferentially along transcriptional states associated with effector differentiation, proliferation, and elevated Ifng expression (Fig. S6).

Given the increased relative representation of the myeloid compartment, which constituted the dominant immune component in induced tumors, we examined tumor-associated myeloid heterogeneity (Fig. 4I and Fig. S7A, B). Resolving myeloid lineages identified dendritic cells, monocytes, neutrophils, mast cells, and macrophages across both conditions (Fig. 4I and Fig. S7A, B). While macrophages constituted the primary myeloid population in both cohorts, subclustering revealed a pronounced redistribution among transcriptionally defined macrophage states (Fig. 4J and Fig. S7C, D). Uninduced CRVV12 (-/-) tumors showed greater representation of Folr2-high, Folr2-low, and Arg1+ TAM states associated with tissue-resident, reparative, and immunoregulatory programs (Fig. 4J, Fig. S7C, D, and Fig. S8A-C)^22,23^. In contrast, dual-switch induction was accompanied by a contraction of these suppressive subsets and the predominant accumulation of immunostimulatory Cxcl9+ and inflammatory Il1b+ TAM populations (Fig. 4J, Fig. S7C, D, and Fig. S8D, E)^23^. These induced macrophage subsets were distinguished by heightened expression of interferon-responsive chemokines (Cxcl9, Cxcl10), antigen-processing machinery, and inflammatory mediators (Il1b, Tnf, Cd40) (Fig. S7C). Together, these analyses associate dual-switch CRVV12 induction with a coordinated shift toward cytotoxic CD8+ T-cell programs and inflammatory Cxcl9+ and Il1b+ macrophage states, revealing qualitative remodeling of the tumor immune microenvironment despite a lower relative lymphoid frequency.

### Model-dependent enhancement of CRVV12 antitumor activity by immune checkpoint blockade

Because dual-switch CRVV12 induction promoted cytotoxic CD8+ T-cell programs and inflammatory macrophage states, we next asked whether immune checkpoint blockade could further amplify its therapeutic activity across five syngeneic tumor models (Fig. 5A). Mice bearing established subcutaneous tumors received intratumoral injections of CRVV12 under dual induction (+Dox +Temsi) alongside systemic administration of anti-PD-1 or anti-CTLA-4 antibodies (Fig. 5A). Therapeutic responses varied markedly across the evaluated tumor models (Fig. 5B, C). In CT26-LacZ tumors, addition of anti-CTLA-4 produced the greatest suppression of tumor outgrowth and the highest long-term survival among all cohorts (Fig. 5B, C). In RENCA tumors, all fully induced CRVV12 cohorts suppressed tumor outgrowth and prolonged survival relative to uninduced virus and PBS controls, with little additional separation conferred by checkpoint inhibitor administration (Fig. 5B, C). In CMT lung carcinoma, fully induced CRVV12 produced modest tumor-growth delay and survival extension, with no clear additional advantage from either checkpoint antibody (Fig. 5B, C). In MC38 colorectal tumors and B16-F10 melanoma, checkpoint combinations were associated with greater tumor-growth control than CRVV12 alone, although the magnitude and survival benefit differed between antibodies and models (Fig. 5B). In these models, anti-CTLA-4 showed a trend toward improved survival in MC38 tumors, whereas anti-PD-1 extended median survival in B16-F10 melanoma (Fig. 5C).

**Fig. 5.**
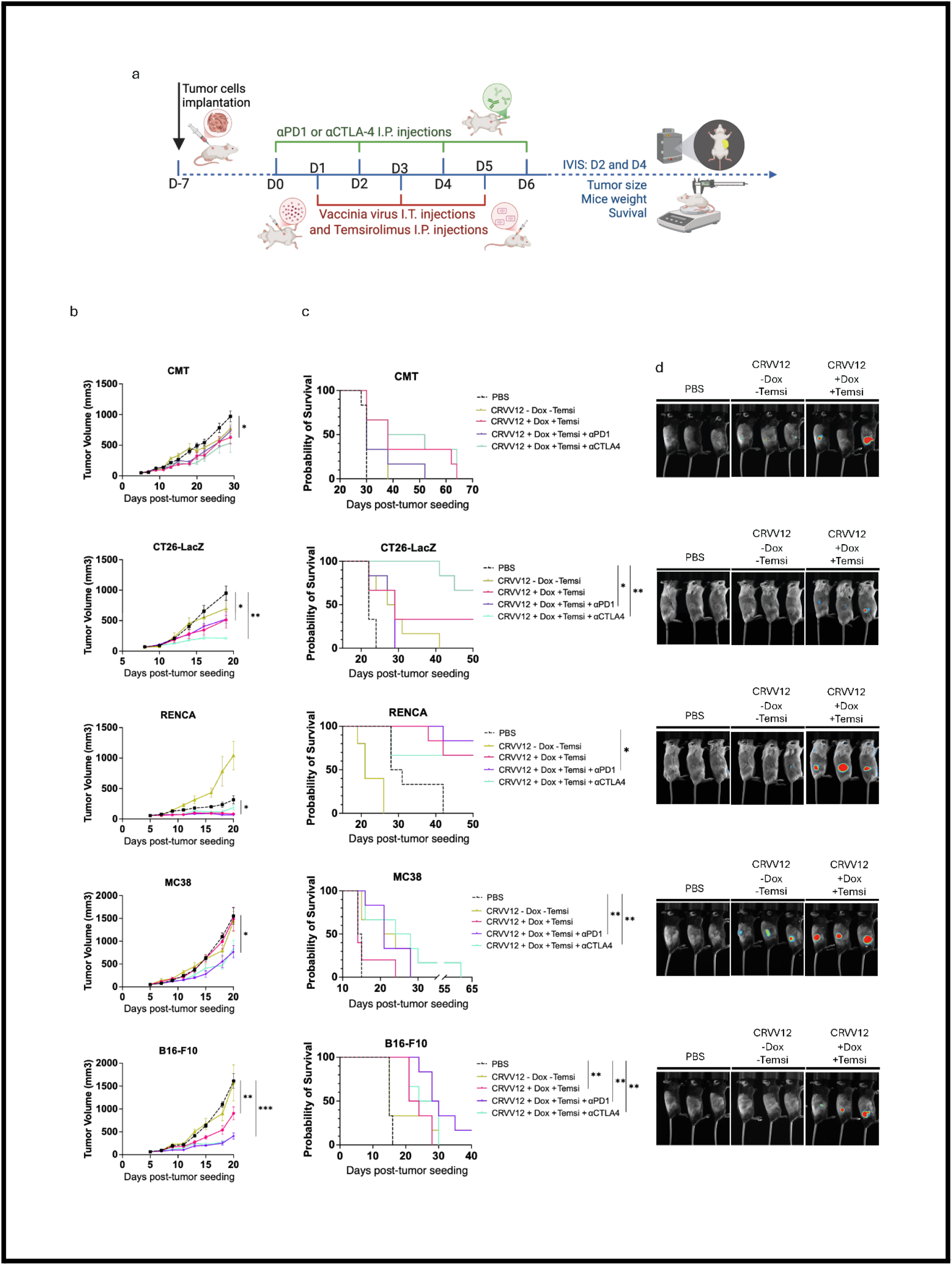
CRVV12 combination with checkpoint blockade produces model-dependent therapeutic responses across syngeneic tumors. (A) Schematic representation of the combination treatment schedule. C57BL/6 mice bearing subcutaneous CMT alveolar lung carcinoma, MC38 colorectal carcinoma, or B16-F10 melanoma, and Balb/c mice bearing subcutaneous CT26 colorectal carcinoma or RENCA renal adenocarcinoma were placed on control or doxycycline diet (625 mg/kg) starting 1 day before treatment initiation. When tumors reached approximately 75 mm³, mice received intraperitoneal injections of isotype control IgG1, anti-PD-1, or anti-CTLA-4 on days 0, 2, 4, and 6, combined with intratumoral injections of PBS or CRVV12 (1 × 10^7 PFU per dose) with intraperitoneal temsirolimus (2.5 mg/kg) or vehicle on days 1, 3, and 5. (B) Longitudinal tumor volume curves for CMT, CT26, RENCA, MC38, and B16-F10 tumor models comparing indicated treatment regimens. (C) Kaplan-Meier survival curves of mice bearing subcutaneous CMT, CT26, RENCA, MC38, and B16-F10 tumors across treatment groups. (D) In vivo bioluminescence imaging (IVIS) of tumor-bearing mice at day 2 post-treatment initiation. In vivo data in (B) represent mean +/- SEM from n = 5 mice per group; P values were determined by two-way ANOVA with Tukey’s post-hoc test. Survival data in (C) represent n = 5 mice per group; P values were determined by log-rank (Mantel-Cox) test. *P < 0.05, P < 0.01, ***P < 0.001, ****P < 0.0001, ns, not significant.

To evaluate viral activity across these distinct tumor environments, we tracked bioluminescent reporter signal at day 2 after treatment initiation (Fig. 5D). Representative imaging demonstrated tumor-associated CRVV12 reporter activity following dual induction across all five models (Fig. 5D). Furthermore, longitudinal monitoring revealed no substantial treatment-associated weight loss in the fully induced CRVV12 combination groups over the monitored interval (Fig. S9). Together, these proof-of-concept evaluations support CRVV12 as a combination platform whose therapeutic response to PD-1 or CTLA-4 blockade varies by tumor context, with the most pronounced additional benefit observed with CTLA-4 blockade in CT26 tumors.

## DISCUSSION

Developing clinically effective oncolytic viruses has long centered on a difficult compromise between replicative potency and host safety^2,3^. Conventional engineering strategies have addressed this challenge primarily through permanent gene deletions, disrupting non-essential metabolic enzymes such as thymidine kinase or viral countermeasures to host innate immunity such as herpes simplex virus 1 ICP34.5^4,5^. While these modifications establish clinically tractable safety profiles and exploit tumor-associated metabolic activity or defective antiviral signaling, they render viral replication contingent on cellular phenotypes that vary widely across and within solid tumors^3,4^. Tumors harbor slow-cycling subclonal populations and microenvironmental niches with intact type I interferon signaling that can restrict the propagation of heavily attenuated vectors^24,25^. At the same time, clinical observations indicate that controlled systemic viral exposure and host immune activation do not invariably lead to unacceptable toxicity. In an exploratory phase I trial evaluating the unattenuated smallpox vaccine strain ACAM2000, viral DNA persisted and reappeared systemically in several patients without inducing severe treatment-associated toxicity. Similarly, in glioblastoma trials evaluating the oncolytic HSV-1 vector CAN-3110, pre-existing HSV-1 seropositivity was associated with longer overall survival and more rapid viral clearance, challenging the assumption that antiviral immunity acts solely as an impediment to therapeutic efficacy^26^. The central engineering challenge is therefore not simply to suppress viral replication universally, but to preserve replication competence within the tumor bed while establishing reliable mechanisms to restrain productive infection after administration.

Resolving this trade-off requires rethinking how viral fitness and selectivity are encoded. Here, we approached this problem through two linked design principles: replacing permanent genetic attenuation with reversible control over essential replication machinery, and complementing entry-based tropism with post-entry control of productive viral persistence. Placing an essential viral gene under dynamic chemogenetic regulation preserves the capacity for robust replication on demand, avoiding permanent fitness penalties that can compromise intratumoral spread. Concurrently, coupling replication to a synchronized, drug-regulated inflammatory payload was designed to shift the tissue environment against productive viral persistence in healthy peripheral organs, and in this study was associated with reduced infectious-virus recovery across normal tissues. Prior platforms have explored cytokine expression to restrict viral virulence, such as the engineering of constitutive interferon-beta expression in thymidine kinase-deleted vaccinia (JX-795)^15^. However, this strategy couples permanent loss of viral functions with immediate inflammatory signaling, which may restrict viral propagation by triggering antiviral clearance before adequate intratumoral expansion. CRVV12 instead places essential replication competence under reversible control while making high-level IL-12 production contingent on an additional chemogenetic input. This hierarchical architecture creates a replication-permissive, payload-restricted state, allowing initial viral propagation to be separated from subsequent inflammatory payload release. The system therefore reduces reliance on permanent attenuation, limits premature immune-mediated restriction of viral spread, and minimizes off-tumor cytokine exposure and the associated risk of systemic inflammatory toxicity.

Translating dynamic replication control to a replicating oncolytic vector required identifying an essential genetic node capable of balancing tight basal repression with robust induced activity. While complete transcriptional shutoff of structural genes such as D13L effectively arrests poxvirus replication for single-cycle vaccine applications^10,18^, our findings show that in the tested D13L architecture, this stringency permitted selection of a TetR frameshift variant during serial in vivo passage. Conversely, the F17R architecture failed to produce meaningful doxycycline-responsive control and acquired loss of TetO sequences during the in vitro escape screen. Targeting E1L, which encodes the catalytic subunit of the viral poly(A) polymerase, VP55, provided the most favorable intermediate phenotype among the loci tested^21^. Uninduced CRVV maintained a restricted basal replication profile that limited systemic viral activity, while doxycycline induction restored viral yields to the range of replication-competent controls. Importantly, this regulatory configuration yielded no detectable consensus-level escape mutations following a single-round in vitro plaque screen or three sequential rounds of in vivo selection. Furthermore, delaying replication induction from day 1 to day 3 post-infection in immunocompromised mice was associated with less weight loss than induction beginning on day 1, supporting temporal control of replication as a means of modulating host tolerability.

Safely deploying interleukin-12 within an oncolytic vector requires stringent suppression of basal expression, as uncontrolled systemic exposure provoked severe systemic inflammatory toxicities in historical clinical trials^17^. By integrating split-T7 RNA polymerase reconstitution with TetR-mediated repression, CRVV12 restricted high-level IL-12 secretion to the fully induced operational state while maintaining low baseline output. Following systemic intravenous administration in athymic CD-1 nude mice, this coordinated circuit produced a striking shift in viral distribution. Whereas control VV-ΔTK and uncoupled CRVV produced widespread extra-tumoral reporter activity, extensive peripheral organ dissemination, and marked morbidity, fully induced CRVV12 maintained stable animal weight and retained high infectious-virus titers within the subcutaneous tumor while sharply reducing recovery from peripheral organs. Infectious virions were below the limit of detection in several major visceral organs and reduced by several orders of magnitude in others, despite sustaining intratumoral titers exceeding 10^5 PFU/g. Because these experiments were conducted in athymic mice, the observed shift in viral biodistribution did not require mature T cells, suggesting that innate immune effectors or local antiviral programs contribute to this altered biodistribution. We propose that coordinated inflammatory payload delivery establishes immune-shaped functional oncotropism after tissue seeding, shifting the balance against productive viral persistence in healthy organs while permitting continued replication within the tumor microenvironment.

High-resolution profiling of treated B16-F10 melanomas revealed that therapeutic efficacy was accompanied by qualitative remodeling of the immune microenvironment despite no increase in relative lymphocyte representation. Dual-switch CRVV12 induction was associated with a lower relative frequency of tumor-associated lymphocytes within the CD45+ compartment and increased relative representation of macrophages. Single-cell transcriptomic profiling showed that this compositional shift was accompanied by transcriptional specialization: tumor-infiltrating CD8+ T cells were preferentially represented within a cytotoxic transcriptional state characterized by higher Ifng expression, while fully induced tumors showed reduced representation of Folr2- and Arg1-expressing macrophage states and greater representation of inflammatory Cxcl9- and Il1b-expressing states^22^. Chemokines such as CXCL9 play a well-established role in guiding effector T-cell positioning and maintaining antitumor activity within the tumor bed^27^. Together, these findings suggest that the quality of the immune states associated with CRVV12 treatment may be more informative than relative lymphocyte abundance alone.

Evaluating CRVV12 in combination with immune checkpoint inhibitors across five syngeneic tumor models demonstrated that therapeutic responses were distinctly context-dependent. While combination therapy produced substantial antitumor activity, the degree of benefit varied considerably across models. The most pronounced enhancement was observed in CT26 colorectal tumors, where the addition of anti-CTLA-4 produced the greatest suppression of tumor outgrowth and the highest long-term survival among the evaluated cohorts. In other models, such as RENCA renal carcinoma, fully induced CRVV12 monotherapy drove robust tumor suppression with little added divergence from checkpoint blockade, while CMT lung carcinoma exhibited modest responses across all viral cohorts. These model-dependent responses likely reflect differences in tumor-intrinsic biology and baseline immune context that were not resolved in the present study. These findings indicate that CRVV12 is unlikely to function as a uniform sensitizer to checkpoint blockade and motivate biomarker-guided selection of combination regimens.

While these findings demonstrate proof of principle for coordinated chemogenetic control and post-entry functional oncotropism, several considerations guide future development. A primary area for further investigation centers on host immunology: because biodistribution and peripheral restriction were evaluated in athymic CD-1 nude mice, establishing CRVV12’s clearance kinetics in the presence of intact adaptive immunity post intravenous injection remains an important next step. In addition, directly quantifying the temporal and spatial exposure of IL-12 across serum and peripheral tissues will further refine the therapeutic window for systemic administration. Finally, expanding combinatorial regimens and testing therapeutic efficacy following intravenous delivery across a broader spectrum of solid tumor models will help identify predictive determinants of response and clarify the full translational potential of this platform.

In conclusion, this work demonstrates that oncolytic vector selectivity does not have to rely exclusively on permanent genetic attenuation or engineered receptor tropism. CRVV12 provides a means to regulate replication competence independently of permanent genomic attenuation while coordinating that control with inflammatory payload output. This architecture preserved high infectious-virus titers in the tumor while producing tumor-biased persistence after systemic delivery, supporting post-entry regulation of productive infection as an additional route to functional oncotropism. Dynamic regulation of viral replication and inflammatory signaling therefore provides an engineering framework for developing potent and manageable oncolytic vectors for heterogeneous solid tumors.

## MATERIALS AND METHODS

### Ethics

Animal procedures were approved by the University of Ottawa Institutional Animal Care Committee (protocols OHRI2870, MEe-2258) and the Animal Protection Committee of the Université de Sherbrooke (protocol 2023-4035) in accordance with Canadian Council on Animal Care guidelines. Biohazardous materials were handled under certificate GC317-125-12.

### Viruses

Conditionally replicating vaccinia viruses were derived from the Tian Tan strain (provided by Dr. David Evans). Inducible replication switches were generated by placing E1L, D13L, or F17R under the control of a synthetic P11 promoter containing a tetracycline operator (TetO), coupled with a PLEO promoter-driven TetR cassette. Constructs co-expressed EGFP and firefly luciferase. For regulated murine IL-12 expression (CRVV12), a TetO-containing T7 promoter driving mIL-12 was integrated at the G1L-I8R locus alongside a split-T7 RNA polymerase fused to FKBP and FRB domains and an mCherry reporter. Control VV-ΔTK was generated by disrupting the thymidine kinase gene (J2R). Transgenes were codon-optimized, designed using SnapGene, and synthesized by GenScript.

### Cell Lines

U2OS, HeLa, Vero, HEK293, HT-29, B16-F10, CT26-LacZ, and MC38 cells were obtained from ATCC and cultured in DMEM or RPMI 1640 supplemented with 10% FBS and 1% penicillin-streptomycin at 37°C in 5% CO2. Co-cultures utilized RPMI supplemented with 10% heat-inactivated FBS. Cell lines were routinely screened for mycoplasma contamination.

### Virus Production and titration

Viruses were propagated in HeLa cells (MOI 0.05, 42 hours), harvested by freeze-thaw lysis, and purified through a 36% sucrose cushion. Viral titers from cell lysates and bead-beaten murine tissue homogenates were determined by plaque assay on U2OS monolayers using an overlay of DMEM and 1% carboxymethyl cellulose. Plaques were scored at 48 hours using an EVOS M5000 imaging system.

### Plaque size assay

Monolayers of HeLa, Vero, or HEK293 cells were infected with VV-TetR-iE1L (MOI 0.0001) for 2 hours, overlaid with carboxymethyl cellulose medium with or without 100 ng/mL doxycycline, and incubated for 96 hours. Plaque areas were imaged using an EVOS M5000 system and quantified using ImageJ.

### Chemokines and cytokines assay

MC38 cells were infected with CRVV or CRVV12 (MOI 0.001) for 48 hours with or without 100 ng/mL doxycycline and 10 nM temsirolimus. Naïve murine splenocytes were isolated following red blood cell lysis and co-cultured with infected tumor cells for 24 hours. Supernatants were subsequently transferred onto fresh naïve splenocytes for secondary incubation. Harvested conditioned media were analyzed using the MD36 Discovery Assay panel by Eve Technologies.

### Western Blot

Cell lysates prepared 18 hours post-infection were resolved by SDS-PAGE and transferred to nitrocellulose membranes. Membranes were probed with primary antibodies against murine IL-12 p35 (R&D Systems), GAPDH (Cell Signaling Technology), or vaccinia virus proteins (LSBio), followed by HRP-conjugated secondary antibodies (Cell Signaling Technology) and chemiluminescent detection (Bio-Rad).

### In Vivo Models

For xenograft models, 1 × 10^6 HT-29 cells in a 1:1 PBS and Matrigel suspension were implanted subcutaneously into female CD-1 nude mice (Charles River Laboratories).

Established tumors (∼50 mm^2) were injected intratumorally with 1 × 10^7 PFU of indicated viruses. For biodistribution studies, mice received 1 × 10^7 PFU intravenously. Doxycycline was administered via formulated diet (625 mg/kg; Bio-Serv), and temsirolimus was delivered intraperitoneally (2.5 mg/kg on days 0, 2, and 4 post-infection). For syngeneic models, C57BL/6J mice (Jackson Laboratory) were injected intraperitoneally with 5 × 10^5 MC38 cells or subcutaneously with 2 × 10^5 B16-F10 cells; BALB/c mice (Charles River Laboratories) received 5 × 10^5 CT26-LacZ cells subcutaneously. Mice received three intratumoral injections of 1 × 10^7 PFU virus. For combination studies, anti-PD-1 (clone RMP1-14; Bio X Cell) or anti-CTLA-4 (clone UC10-4F10-11; Bio X Cell) was administered intraperitoneally at 5 mg/kg for four doses. Viral bioluminescence was imaged 5 minutes after intraperitoneal injection of D-luciferin (2 mg/mouse; MedChemExpress). Tumor dimensions and animal weights were monitored longitudinally.

### Flow Cytometry and Phenotyping

Tumors were dissociated using the Mouse Tumor Dissociation Kit (Miltenyi Biotec), and single-cell suspensions from tumors and spleens were stained with fluorochrome-conjugated antibodies. Samples were acquired on a BD LSRFortessa cytometer and analyzed using FlowJo.

### Single-cell RNA sequencing and bioinformatic analysis

Sequencing libraries were prepared using 10x Genomics protocols. FASTQ files were aligned to the GRCm39 reference genome using Cell Ranger and analyzed in Seurat (v5.4.0). Cells expressing 200–7,500 genes with 500–40,000 UMIs and <10% mitochondrial reads were retained, and doublets were filtered using DoubletFinder (v2.0.6). Data were log-normalized, scaled regressing UMI depth, and integrated across conditions using Harmony (v1.2.4) across 20 principal components (clustering resolution 0.8). Macrophage trajectories were constructed in Monocle 3 (v1.4.26).

### Statistical Analysis

Analyses were performed in GraphPad Prism using two-tailed unpaired Student’s t-tests, one-way ANOVA with post-hoc tests, or log-rank tests for survival as specified in figure legends. Statistical significance was defined as P < 0.05 (*P < 0.0332; P < 0.0021; ***P < 0.0002; ****P < 0.0001).

## Author contributions

We thank Kathy Sheikheleslamy and the Ottawa Hospital Research Institute StemCore Laboratories for assistance with single-cell RNA sequencing, and Fernando Ortiz and the Ottawa Hospital Research Institute Flow Cytometry and Cell Sorting Facility for assistance with flow cytometry and cell sorting.

## Declaration of Generative AI and AI-assisted technologies in the manuscript preparation process

During the preparation of this work, the authors used ChatGPT and Google Gemini to assist with language editing, improving readability, and polishing manuscript phrasing. The authors critically reviewed, revised, and edited all outputs and take full responsibility for the final content of the published article.

---

**Fig. S1.**
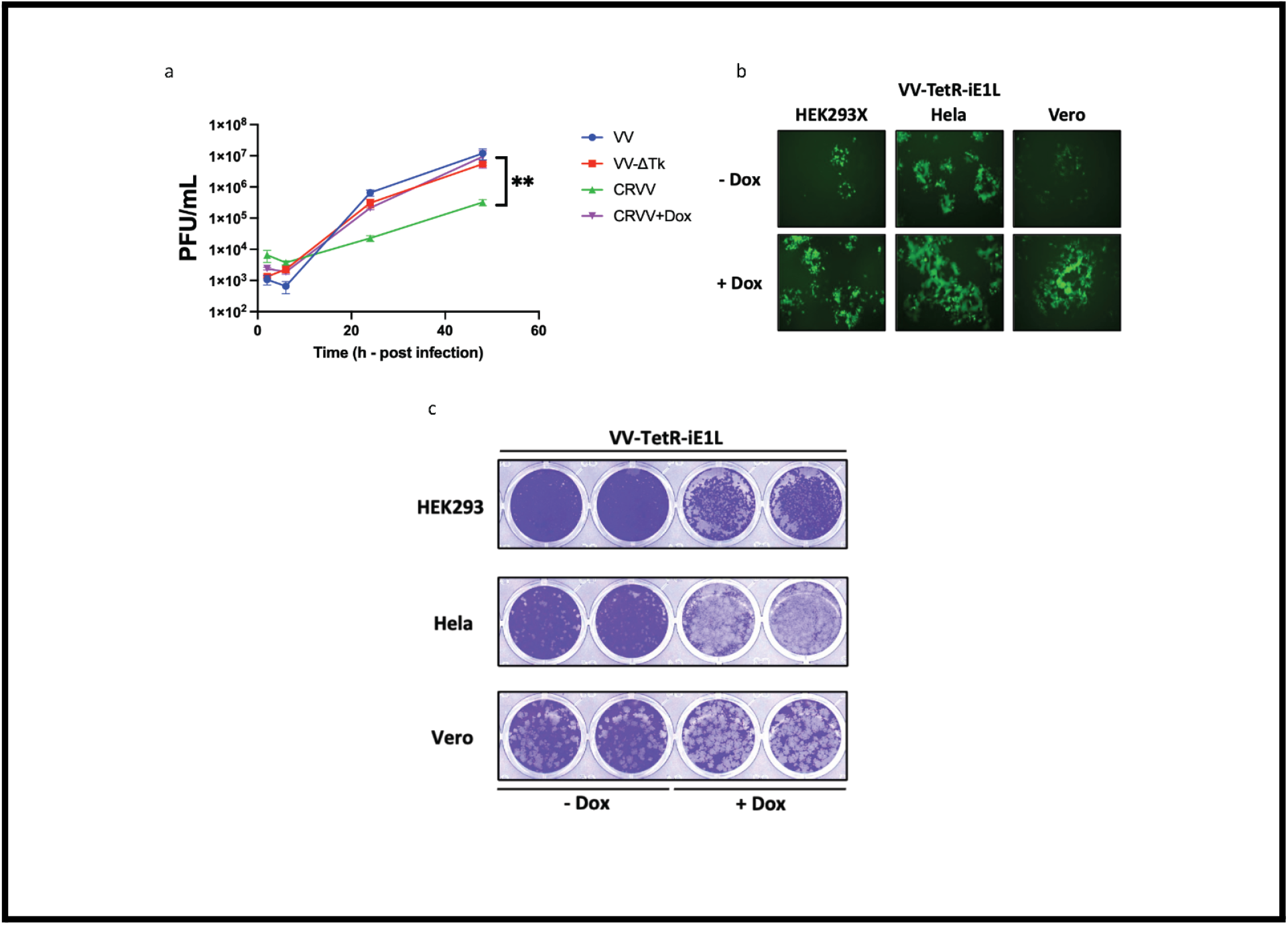
Stringent repression of E1L restricts viral replication and plaque formation across multiple cell lines while preserving wild-type kinetics upon induction. (A) Multi-step viral growth curves in HeLa cells infected with wild-type VV, VV-ΔTK, or CRVV (MOI 0.01) in the absence (-Dox) or presence (+Dox; 100 ng/mL) of doxycycline. Total viral progeny were harvested at 2, 6, 24, and 48 h post-infection and titered on U2OS cells. (B) Representative fluorescence microscopy images of HEK293, HeLa, and Vero cells infected with VV-TetR-iE1L (MOI 0.01) at 48 h post-infection with or without doxycycline (100 ng/mL) in the media. (C) Crystal violet plaque and cytopathic effect staining of HEK293, HeLa, and Vero cell monolayers infected with VV-TetR-iE1L (MOI 0.0001) in the absence (-Dox) or presence (+Dox; 100 ng/mL) of doxycycline at 96 h post-infection. Data in (A) represent mean +/- SEM from n = 3 replicates. P values were determined by two-way ANOVA. P < 0.01.

**Fig. S2.**
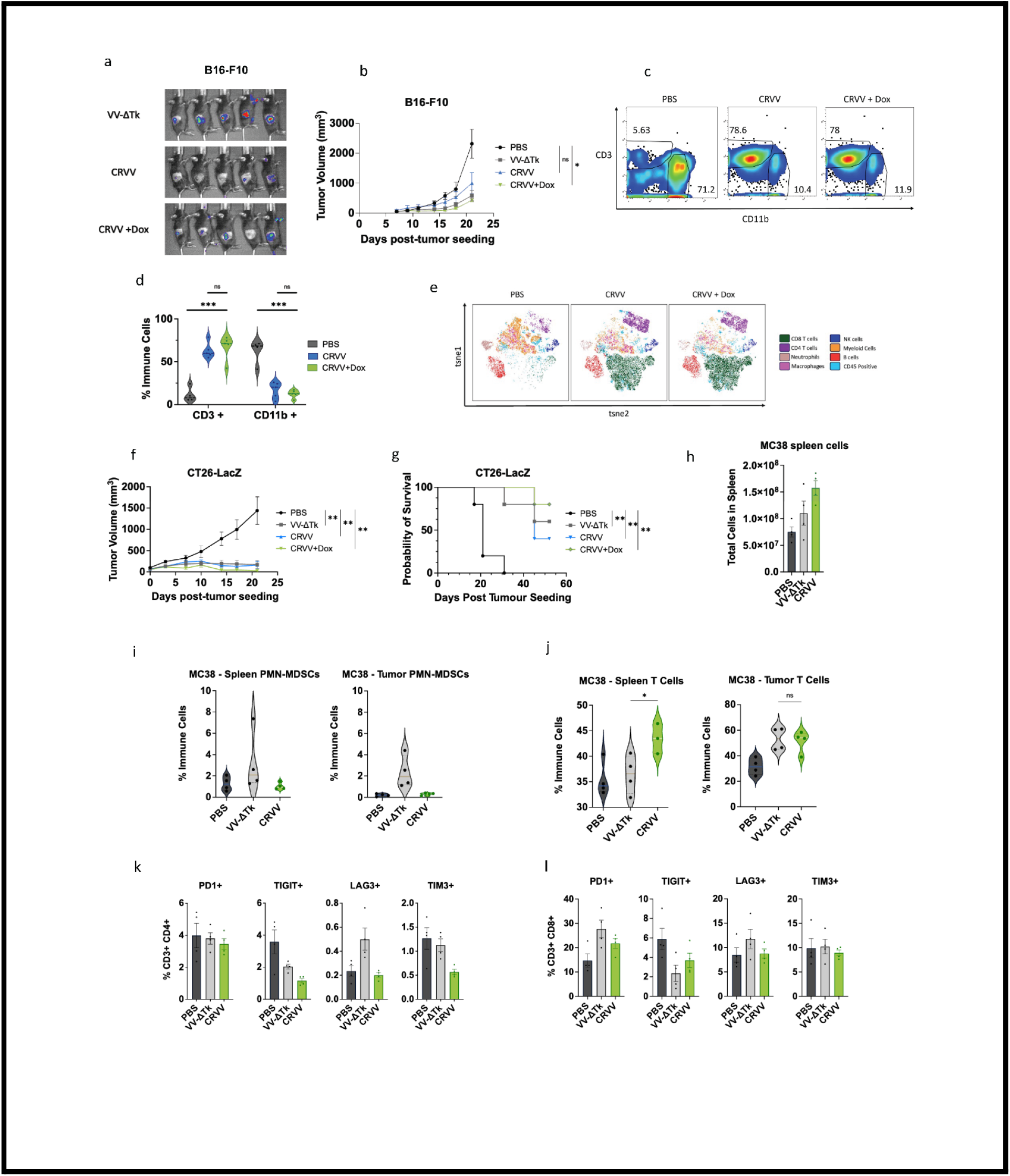
CRVV promotes systemic T cell expansion and drives distinct immune remodeling in B16-F10 and MC38 tumor models. (A) In vivo bioluminescence imaging (IVIS) of C57BL/6 mice bearing subcutaneous B16-F10 melanoma tumors at day 5 post-intratumoral injection of 1 × 10^7 PFU VV-ΔTK, CRVV, or CRVV on a doxycycline-supplemented diet. (B) Subcutaneous B16-F10 tumor volume trajectories following intratumoral injection of VV-ΔTK or CRVV (1 × 10^7 PFU) with or without doxycycline diet. (C) Representative flow cytometry plots of CD3 and CD11b expression on CD45+ tumor-infiltrating immune cells isolated from subcutaneous B16-F10 tumors 7 days post-intratumoral injection of PBS, CRVV (-Dox), or CRVV (+Dox diet). (D) Violin plots showing the percentage of CD3+ and CD11b+ immune cells within B16-F10 tumors across indicated treatment groups. (E) t-SNE visualization of flow cytometrically profiled tumor-infiltrating CD45+ immune cell subsets in B16-F10 tumors following treatment with PBS, CRVV, or CRVV + Dox. (F) Longitudinal tumor volume measurements in Balb/c mice bearing subcutaneous CT26-LacZ tumors treated with intratumoral injections of PBS, VV-ΔTK, CRVV, or CRVV + Dox (1 × 10^7 PFU). (G) Kaplan-Meier survival curves of Balb/c mice bearing CT26-LacZ tumors tracked through day 60 post-tumor inoculation. (H) Absolute cell counts of total splenocytes harvested from C57BL/6 mice bearing subcutaneous MC38 tumors 6 days post-intratumoral injection of PBS, VV-ΔTK, or CRVV (1 × 10^7 PFU) on a doxycycline-supplemented diet. (I) Violin plots quantifying the percentage of PMN-MDSCs in the spleen (left) and tumor (right) of MC38 tumor-bearing mice 6 days post-treatment. (J) Violin plots quantifying the percentage of total T cells in the spleen (left) and tumor (right) across treatment groups in MC38 tumor-bearing mice. (K) Percentage of tumor-infiltrating CD3+ CD4+ T cells (left) and CD3+ CD8+ T cells (right) expressing PD-1, TIGIT, LAG-3, and TIM-3 at 6 days post-treatment. Data represent mean +/- SEM from n = 4 or 5 mice per group. Statistical significance was determined by two-way ANOVA for tumor volumes (B, F), log-rank test for survival (G), and one-way ANOVA with Tukey’s post-hoc test for flow cytometric and cell count data (D, H, K). *P < 0.05, P < 0.01, ***P < 0.001, ns, not significant.

**Fig. S3.**
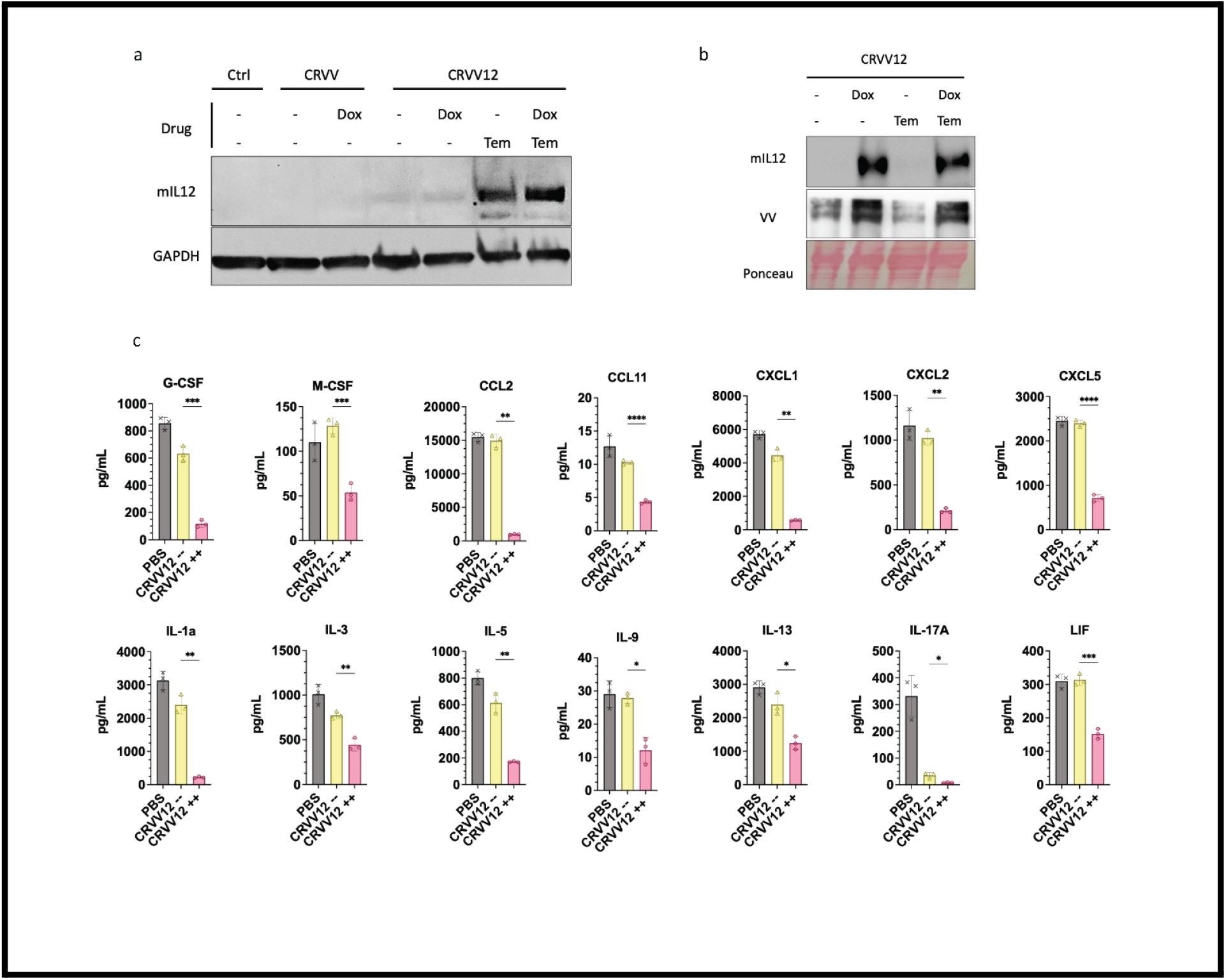
Validation of dual-switch interleukin-12 regulation and extended cytokine profiling following splenocyte coculture. (mIL-12) protein levels in cell lysates at 18 h post-infection with control, CRVV, or CRVV12 (MOI 0.5) in the absence or presence of doxycycline (100 ng/mL), temsirolimus (10 nM), or both. GAPDH was used as a loading control. (B) Western blot analysis of mIL-12 protein levels in lysates of U2OS cells infected with CRVV12 (MOI 0.01) at 48 h post-infection under the indicated drug conditions. Ponceau S total protein staining was used as a loading control. (C) Absolute concentrations (pg/mL) of G-CSF, M-CSF, CCL2, CCL11, CXCL1, CXCL2, CXCL5, IL-1α, IL-3, IL-5, IL-9, IL-13, IL-17A, and LIF released from naive Balb/c splenocytes co-cultured for 24 h with filtered conditioned media from MC38 cells infected with CRVV12 (MOI 0.001) under uninduced (CRVV12 -/-) or fully induced (CRVV12 +/+) conditions relative to PBS controls. Data in (C) represent mean +/- SEM from n = 3 replicates. P values were determined by one-way ANOVA with Tukey’s post-hoc test. *P < 0.05, P < 0.01, ***P < 0.001, ****P < 0.0001.

**Fig. S4.**
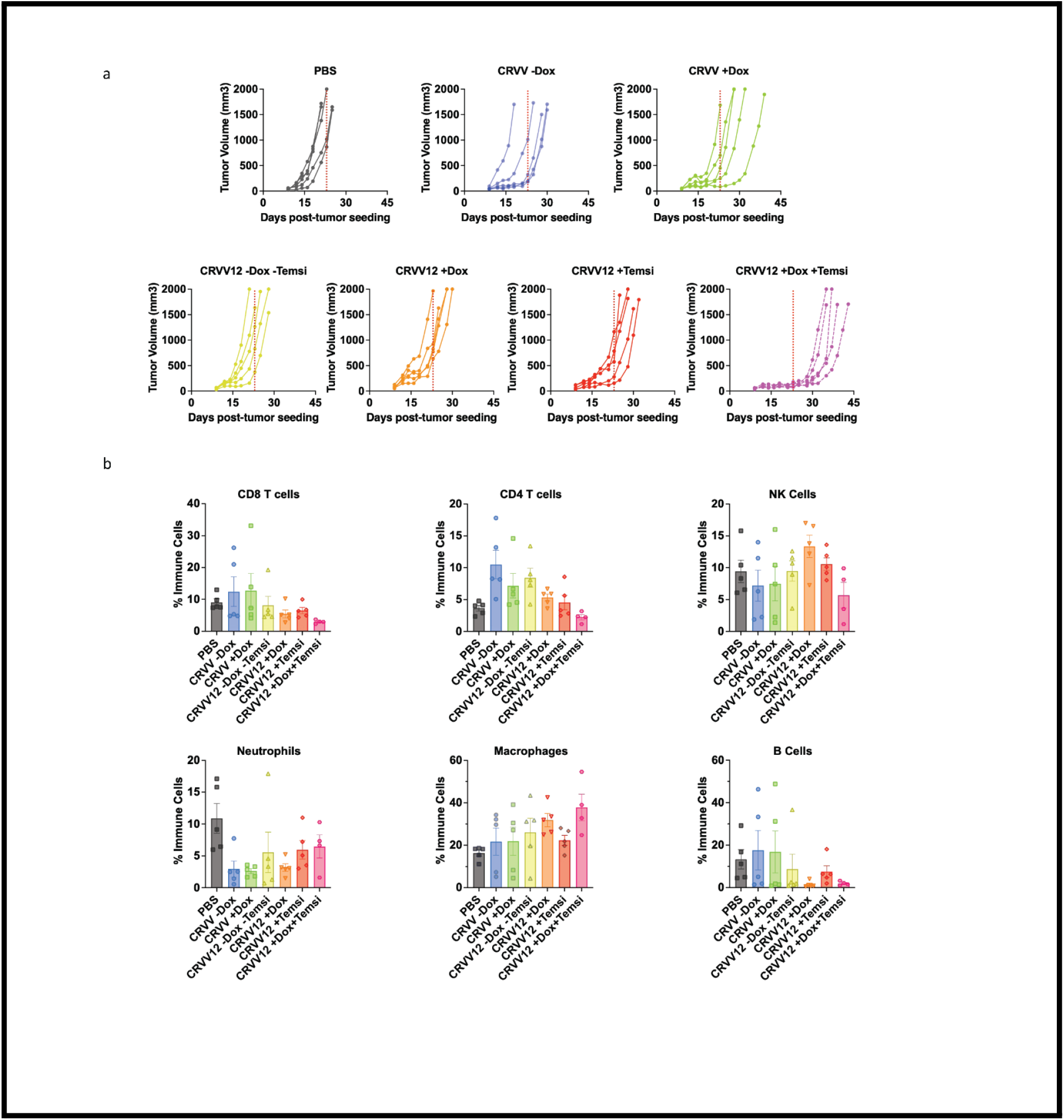
Individual tumor growth trajectories and flow cytometric quantification of tumor-infiltrating immune subsets across CRVV12 operational states. (A) Individual subcutaneous B16-F10 melanoma tumor growth curves over time in C57BL/6 mice treated with intratumoral injections of PBS, CRVV -Dox, CRVV +Dox, CRVV12 -Dox -Temsi, CRVV12 +Dox, CRVV12 +Temsi, or CRVV12 +Dox +Temsi (1 × 10^7 PFU per dose). (B) Flow cytometric quantification of major tumor-infiltrating immune subsets (percentage of total CD45+ leukocytes) harvested 4 days post-injection across indicated treatment groups. Data in (A) and (B) represent mean +/- SEM from n = 5 mice per group. P values in (B) were determined by one-way ANOVA with Tukey’s post-hoc test. *P < 0.05, P < 0.01, ***P < 0.001, ns, not significant .

**Fig. S5.**
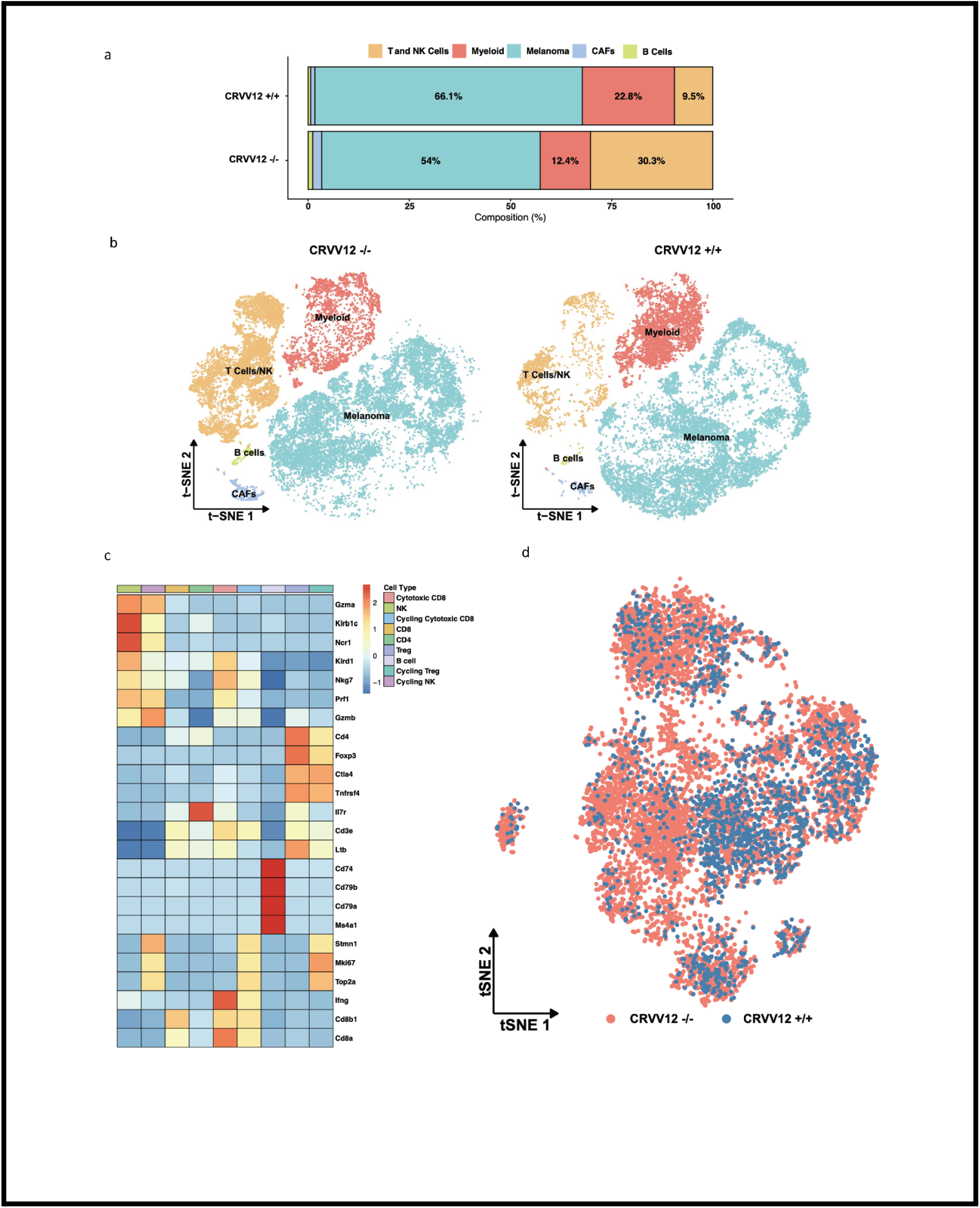
Global single-cell transcriptomic profiling and marker-based annotation of tumor-infiltrating lymphoid subpopulations. (A) Stacked bar chart displaying overall cellular composition (%) of B16-F10 tumors in CRVV12 +/+ versus CRVV12 -/- conditions. (B) Global t-SNE projections of annotated cell lineages split by CRVV12 -/- and CRVV12 +/+ conditions. (C) Heatmap showing row-normalized expression of canonical lineage-defining markers across annotated lymphoid subsets. (D) t-SNE projection of the lymphoid compartment colored by treatment condition (CRVV12 -/- vs CRVV12 +/+).

**Fig. S6.**
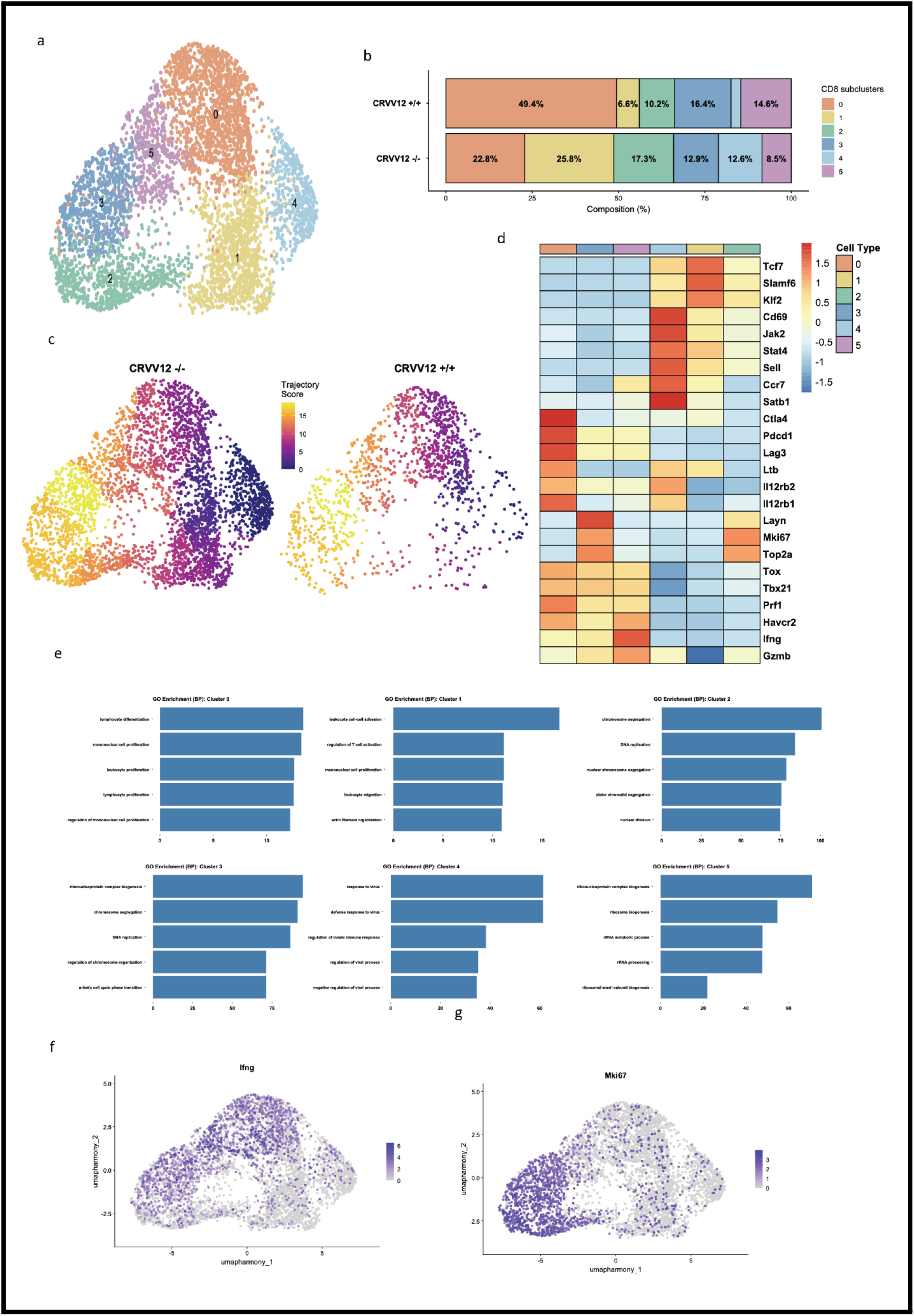
Pseudotime analysis associates CRVV12 dual induction with proliferative and IFNγ-expressing CD8+ T-cell states. (A) UMAP projection of the CD8+ T cell compartment resolving subclusters 0 to 5. (B) Stacked bar chart showing the relative proportion (%) of CD8+ T cell subclusters in CRVV12 +/+ versus CRVV12 -/- tumors. (C) Pseudotime differentiation trajectory score mapped onto UMAP projections of CD8+ T cells split by treatment condition. (D) Heatmap of row-normalized expression for selected stem-like, activation, proliferation, exhaustion, and cytotoxic effector genes across CD8+ T cell clusters 0 to 5. (E) Gene Ontology (GO) biological process enrichment analysis for CD8+ T cell clusters 0 through 5. (F) UMAP feature plot of normalized Ifng expression in CD8+ T cells. (G) UMAP feature plot of normalized Mki67 expression in CD8+ T cells.

**Fig. S7.**
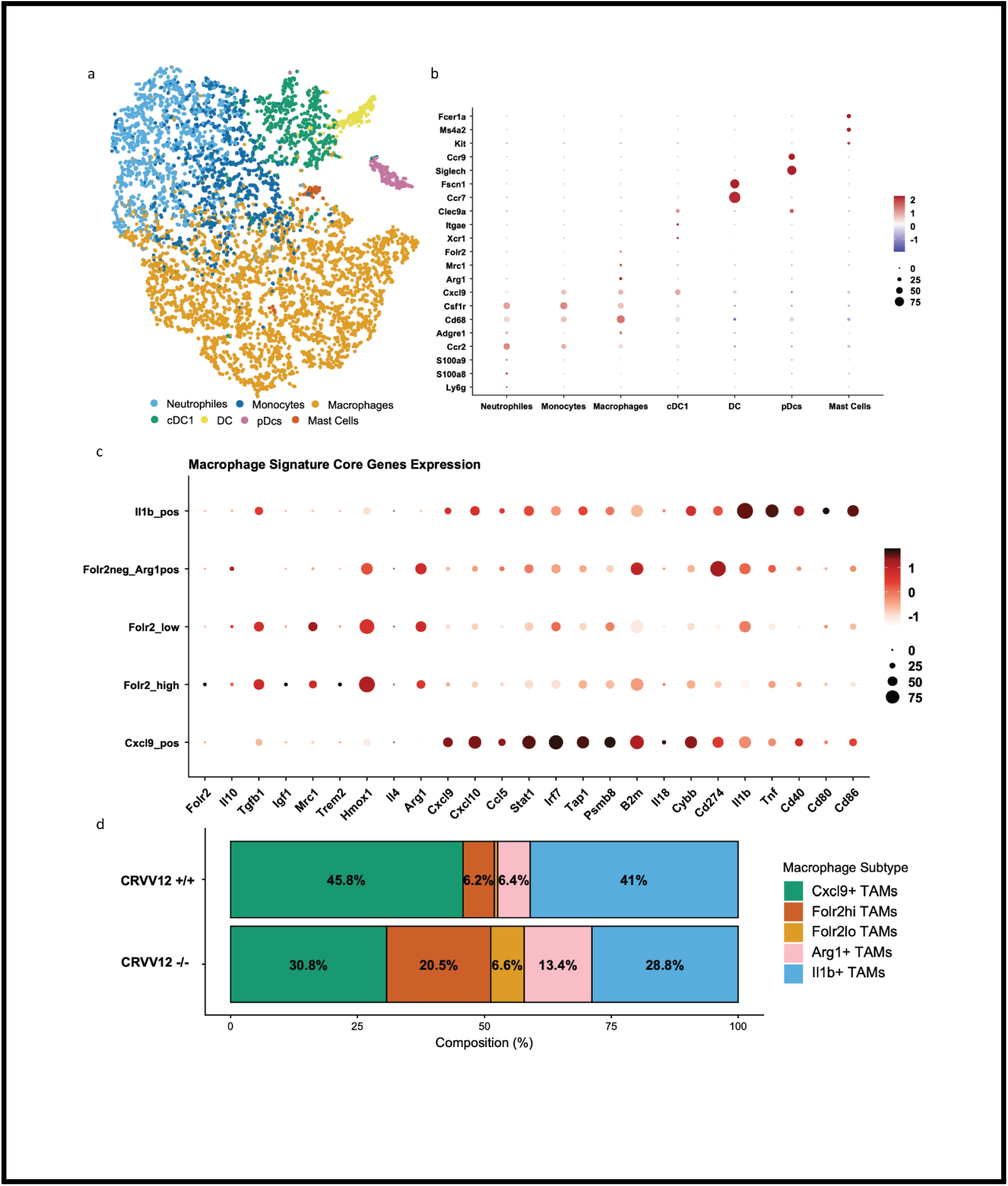
Transcriptomic marker profiling and compositional quantification of tumor-associated myeloid lineages and macrophage subsets. (A) UMAP projection of tumor-infiltrating myeloid cells resolving major myeloid lineages. (B) Dot plot showing expression level and percentage of expressing cells for lineage-defining genes across myeloid subsets. (C) Dot plot displaying expression of core signature genes defining immunosuppressive, tissue-resident, inflammatory, and antigen-presenting programs across macrophage subtypes. (D) Stacked bar chart showing relative composition (%) of macrophage subtypes in CRVV12 +/+ versus CRVV12 -/- tumors.

**Fig. S8.**
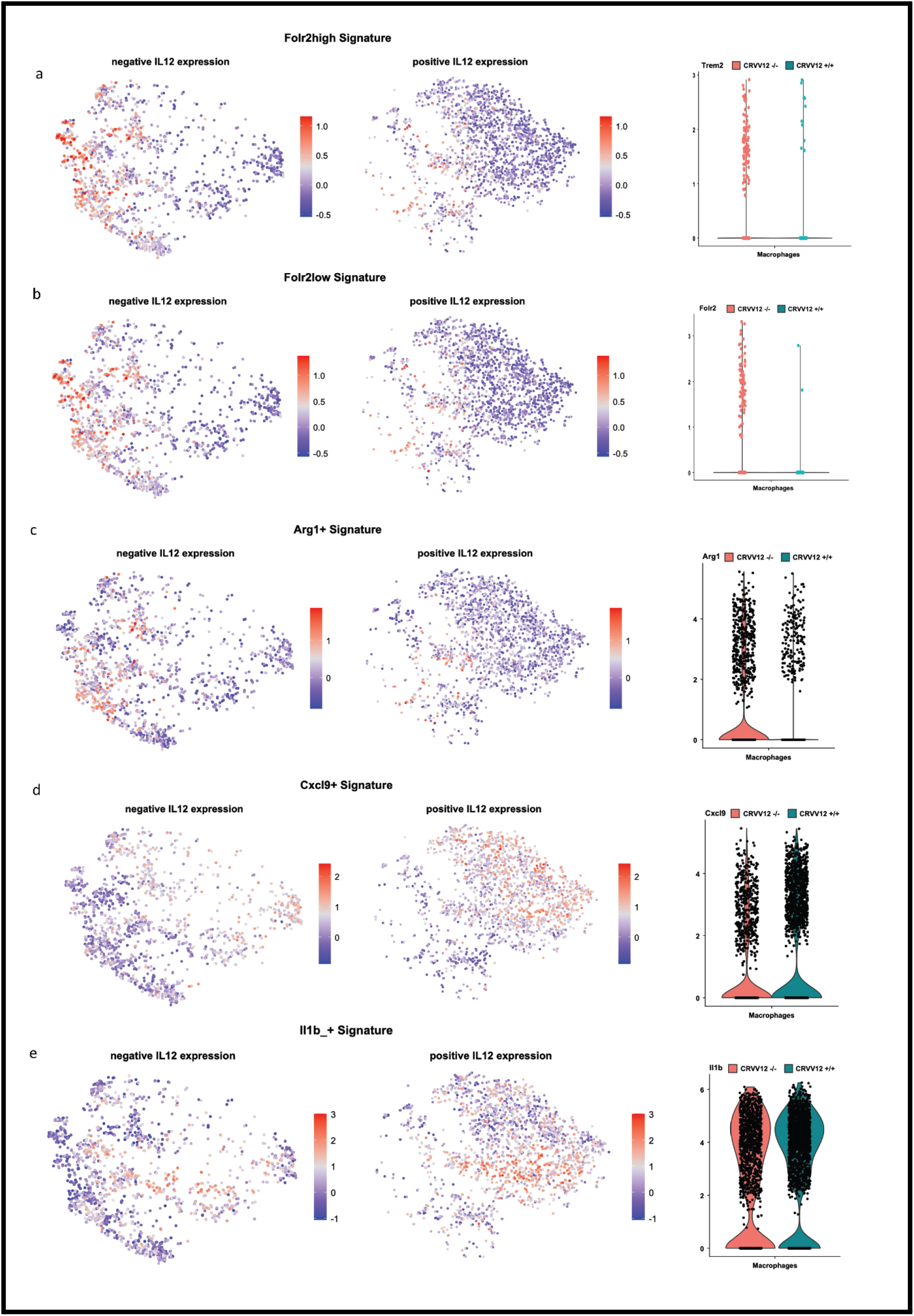
Single-cell expression profiles of core signature genes defining immunosuppressive, tissue-resident, and inflammatory macrophage subsets. (A) Folr2high signature score distribution and violin plots of Trem2 and Folr2 expression in macrophages from CRVV12 -/- and CRVV12 +/+ tumors. (B) Folr2low signature score distribution in macrophages from CRVV12 -/- and CRVV12 +/+ tumors. (C) Arg1+ signature score distribution and violin plot of Arg1 expression in macrophages from CRVV12 -/- and CRVV12 +/+ tumors. (D) Cxcl9+ signature score distribution and violin plot of Cxcl9 expression in macrophages from CRVV12 -/- and CRVV12 +/+ tumors. (E) Il1b+ signature score distribution and violin plot of Il1b expression in macrophages from CRVV12 -/- and CRVV12 +/+ tumors.

**Fig. S9.**
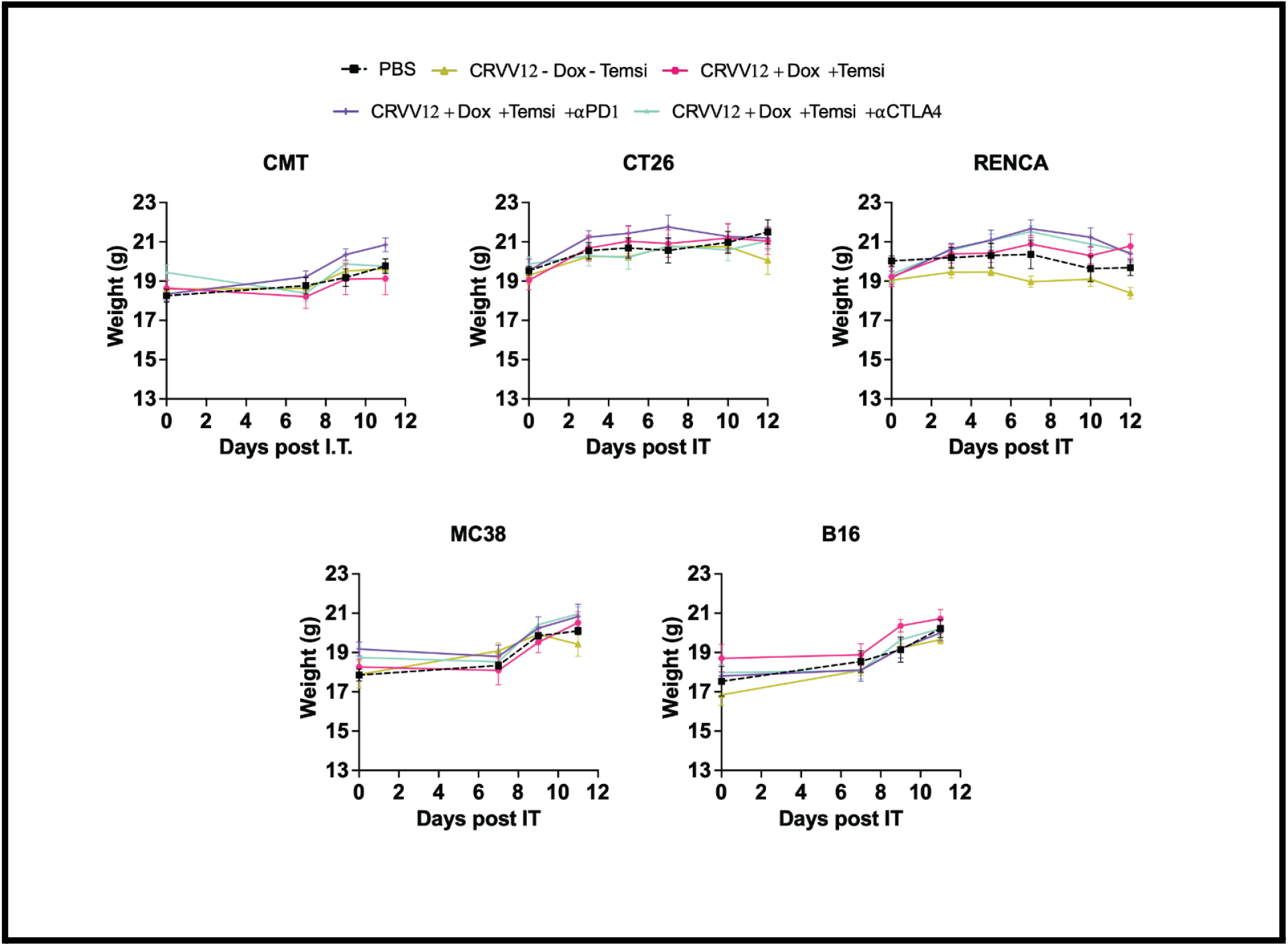
Longitudinal body weight monitoring confirms the systemic safety of CRVV12 combination therapy with immune checkpoint blockade across multiple tumor models. Longitudinal body weight tracking of C57BL/6 mice (bearing subcutaneous CMT, MC38, or B16-F10 tumors) and Balb/c mice (bearing subcutaneous CT26 or RENCA tumors) treated with PBS or CRVV12 (1 × 10^7 PFU) in combination with isotype control IgG1, anti-PD-1, or anti-CTLA-4 antibodies under the indicated drug regimens. Data represent mean +/- SEM from n = 5 mice per group.

